# A glycan gate controls opening of the SARS-CoV-2 spike protein

**DOI:** 10.1101/2021.02.15.431212

**Authors:** Terra Sztain, Surl-Hee Ahn, Anthony T. Bogetti, Lorenzo Casalino, Jory A. Goldsmith, Evan Seitz, Ryan S. McCool, Fiona L. Kearns, Francisco Acosta-Reyes, Suvrajit Maji, Ghoncheh Mashayekhi, J. Andrew McCammon, Abbas Ourmazd, Joachim Frank, Jason S. McLellan, Lillian T. Chong, Rommie E. Amaro

**Affiliations:** Department of Chemistry and Biochemistry, UC San Diego, La Jolla, CA 92093; Department of Chemistry, University of Pittsburgh, Pittsburgh, PA 15260; Department of Molecular Biosciences, The University of Texas at Austin, Austin, TX 78712; Department of Biological Sciences, Columbia University, New York, NY, 10032, USA; Department of Biochemistry and Molecular Biophysics, Columbia University Medical Center, New York, NY 10032, USA; Department of Physics, University of Wisconsin-Milwaukee, 3135 N. Maryland Ave, Milwaukee, WI 53211, USA; Department of Pharmacology, UC San Diego, La Jolla, CA 92093

## Abstract

SARS-CoV-2 infection is controlled by the opening of the spike protein receptor binding domain (RBD), which transitions from a glycan-shielded “down” to an exposed “up” state in order to bind the human ACE2 receptor and infect cells. While snapshots of the “up” and “down” states have been obtained by cryoEM and cryoET, details of the RBD opening transition evade experimental characterization. Here, over 130 μs of weighted ensemble (WE) simulations of the fully glycosylated spike ectodomain allow us to characterize more than 300 continuous, kinetically unbiased RBD opening pathways. Together with ManifoldEM analysis of cryo-EM data and biolayer interferometry experiments, we reveal a gating role for the N-glycan at position N343, which facilitates RBD opening. Residues D405, R408, and D427 also participate. The atomic-level characterization of the glycosylated spike activation mechanism provided herein achieves a new high-water mark for ensemble pathway simulations and offers a foundation for understanding the fundamental mechanisms of SARS-CoV-2 viral entry and infection.

## Main Text

### Introduction

Severe acute respiratory syndrome coronavirus 2 (SARS-CoV-2) is an enveloped RNA virus and the causative agent of COVID-19, a disease that has caused significant morbidity and mortality worldwide.^1,2^ The main infection machinery of the virus is the spike (S) protein, which sits on the outside of the virus, is the first point of contact that the virion makes with the host cell, and is a major viral antigen.^3^ A significant number of cryoEM structures of the spike protein have been recently reported, collectively informing on structural states of the spike protein. The vast majority of resolved structures fall into either “down” or “up” states, as defined by the position of the receptor binding domain (RBD), which modulates interaction with the ACE2 receptor for cell entry.^4,5,6^

The RBDs must transition from a “down” to an “up” state for the receptor binding motif to be accessible for ACE2 binding (**Fig. 1**), and therefore the activation mechanism is essential for cell entry. Mothes et al.^7^ used single-molecule fluorescence (Förster) resonance energy transfer (smFRET) imaging to characterize spike dynamics in real-time. Their work showed that the spike dynamically visits four distinct conformational states, the population of which are modulated by the presence of the human ACE2 receptor and antibodies. However, smFRET, as well as conventional structural biology techniques, are unable to inform on the atomic-level mechanisms underpinning such dynamical transitions. Recently, all-atom molecular dynamics (MD) simulations of the spike protein with experimentally accurate glycosylation together with corroborating experiments indicated the extensive shielding by spike glycans, as well as a mechanical role for glycans at positions N165 and N234 in supporting the RBD in the “open” conformation.^8^ Conventional MD simulations as performed in Casalino et al.^8^ also revealed microsecond-timescale dynamics to better characterize the spike dynamics but were limited to sampling configurations that were similar in energy to the cryoEM structures. Several enhanced sampling MD simulations have been performed to study this pathway, however, these simulations lacked glycosylation for the spike protein^9^ or involve the addition of an external force^10^ or did not provide mechanistic detail.^11^

**Figure 1.**
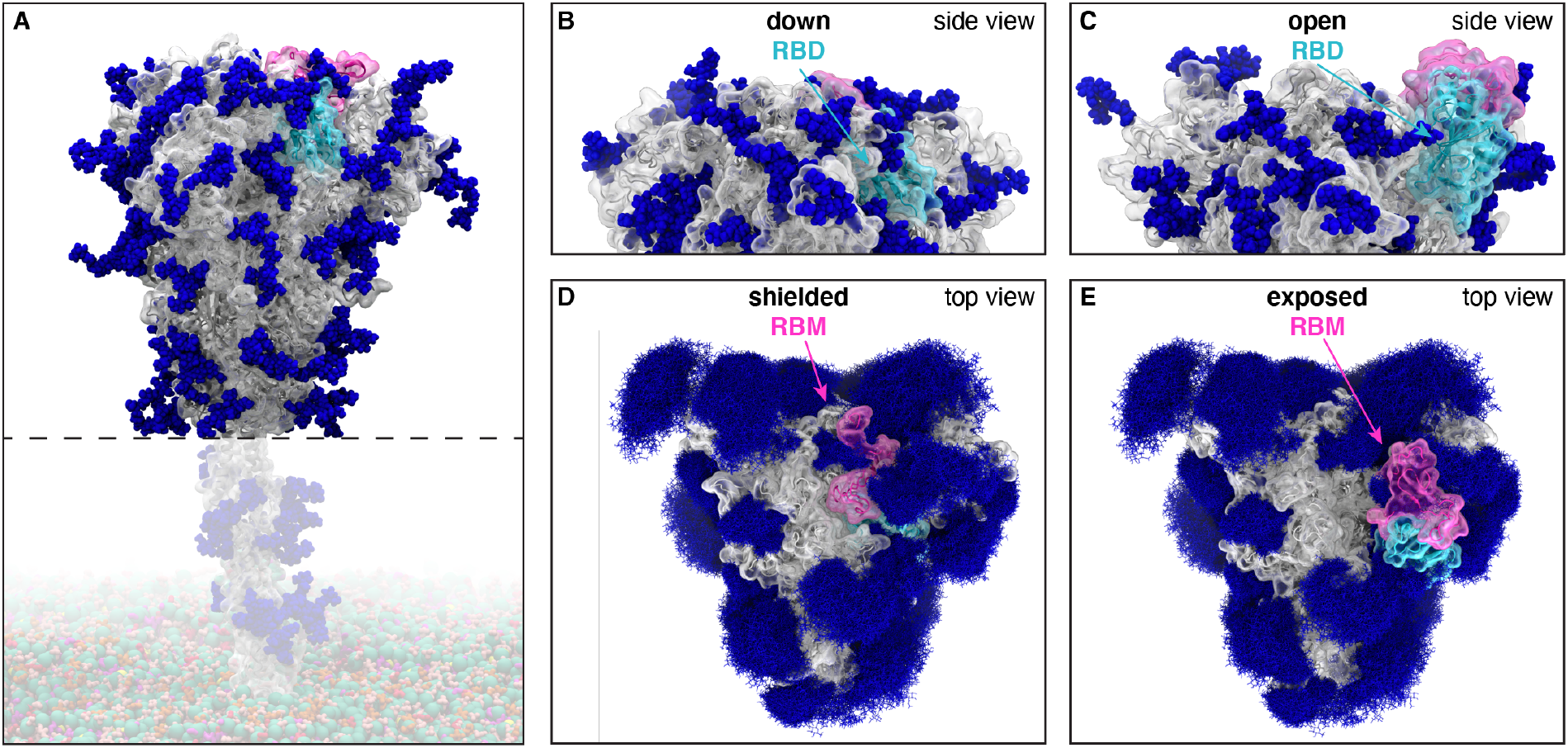
Glycosylated spike RBD “down” and “open” conformations. (A) The SARS-CoV-2 spike head (gray) with glycans (dark blue) as simulated, with the stalk domain and membrane (not simulated here, but shown in transparent for completeness). RBD shown in cyan, receptor binding motif (RBM) in pink. Side view of the “down” (shielded, B) and “open” (exposed, C) RBD. Top view of the closed (shielded, D) and “open” (exposed, E) RBM. Composite image of glycans (dark blue lines) shows many overlapping snapshots of the glycans over the microsecond simulations.

In this study, we characterized the spike RBD opening pathway for the fully glycosylated SARS-CoV-2 spike protein, in order to gain a detailed understanding of the activation mechanism. We used the weighted ensemble (WE) path sampling strategy^12,13^ (**Supplementary Fig. 1**) to enable the simulation of atomistic pathways for the spike opening process. As a path sampling strategy, WE focuses computing power on the functional transitions between stable states rather than the stable states themselves.^14^ This is achieved by running multiple trajectories in parallel and periodically replicating trajectories that have transitioned from previously visited to newly visited regions of configurational space,^15^ thus minimizing the time spent waiting in the initial stable state for “lucky” transitions over the free energy barrier. Given that these transitions are much faster than the waiting times,^16,17^ the WE strategy can be orders of magnitude more efficient than conventional MD simulations in generating pathways for rare events such as protein folding and protein binding.^18,19^ This efficiency is even higher for slower processes, increasing *exponentially* with the effective free energy barrier.^20^ Not only are dynamics carried out without any biasing force or modifications to the free energy landscape, but suitable assignment of statistical weights to trajectories provides an *unbiased* characterization of the system’s time-dependent ensemble properties.^13^ The WE strategy therefore generates continuous pathways with unbiased dynamics, yielding the most direct, atomistic views for analyzing the mechanism of functional transitions, including elucidation of transient states that are too fleeting to be captured by laboratory experiments. Furthermore, while the strategy requires a progress coordinate toward the target state, the definition of this target state need not be fixed in advance when applied under equilibrium conditions,^21^ enabling us to refine the definition of the target “open” state of the spike protein based on the probability distribution of protein conformations sampled by the simulation.

Our work characterizes a series of transition pathways of the spike opening, in agreement with conformations detected in cryoEM dataset by ManifoldEM,^22^ and identifies key residues, including a glycan at position N343, that participate in the opening mechanism. Our simulation findings are corroborated by biolayer interferometry experiments, which demonstrate a reduction in the ability of the spike to interact with ACE2 after mutation of these key residues.

## Results and Discussion

### Weighted ensemble simulations of spike opening

As mentioned above, simulations of the spike opening process require an enhanced sampling strategy as the process occurs beyond the microsecond timescale (*i.e.* seconds timescale^7^). We therefore used the weighted ensemble (WE) path sampling strategy, which enabled the generation of continuous, atomistic pathways for the spike opening process with unbiased dynamics (**Fig. 2A-E, Supplementary Video 1**); these pathways were hundreds of ns long, excluding the waiting times in the initial “down” state. The protein model was based on the head region (residues 16 to 1140) of the glycosylated SARS-CoV-2 spike from Casalino et al.^8^ (**Fig. 1**), which in turn was built on the cryoEM structure of the three-RBD-down spike (PDB ID: 6VXX^5^). The entire simulation system, including explicit water and salt ions, reaches almost half a million atoms. We focused sampling along a two-dimensional progress coordinate to track RBD opening: the difference in the center of mass of the spike core to the RBD and the root mean squared deviation (RMSD) of the RBD (**Fig. 2F-G**). On the SDSC Comet and TACC Longhorn supercomputers, 100 GPUs ran the WE simulations in parallel for over a month, generating over 130 μs of glycosylated spike trajectories and more than 200 TB of trajectory data. We simulated a total of 310 independent pathways, including 204 pathways from the RBD-down conformation (PDB ID: 6VXX^5^) to the RBD-up conformation (PDB ID: 6VSB^4^) and 106 pathways from the RBD-down to the RBD-open state, in which the RBD twists open beyond the 6VSB^4^ cryoEM structure. Remarkably, the RBD-open state that we sampled includes conformations that align closely with the ACE2-bound spike cryoEM structure (PDB ID: 7A95^6^) even though this structure was not a target state of our progress coordinate (**Fig. 2F-G**, **Supplementary Video 1, Supplementary Figs. 2 and 3**). This result underscores the value of using (i) equilibrium WE simulations that do not require a fixed definition of the target state and (ii) a two-dimensional progress coordinate that allows the simulations to sample unexpected conformational space along multiple degrees of freedom. The ACE2-bound spike conformation has also been sampled by the Folding@home distributed computing project,^11^ and RBD rotation has been detected in cryoEM experiments.^6^

**Figure 2.**
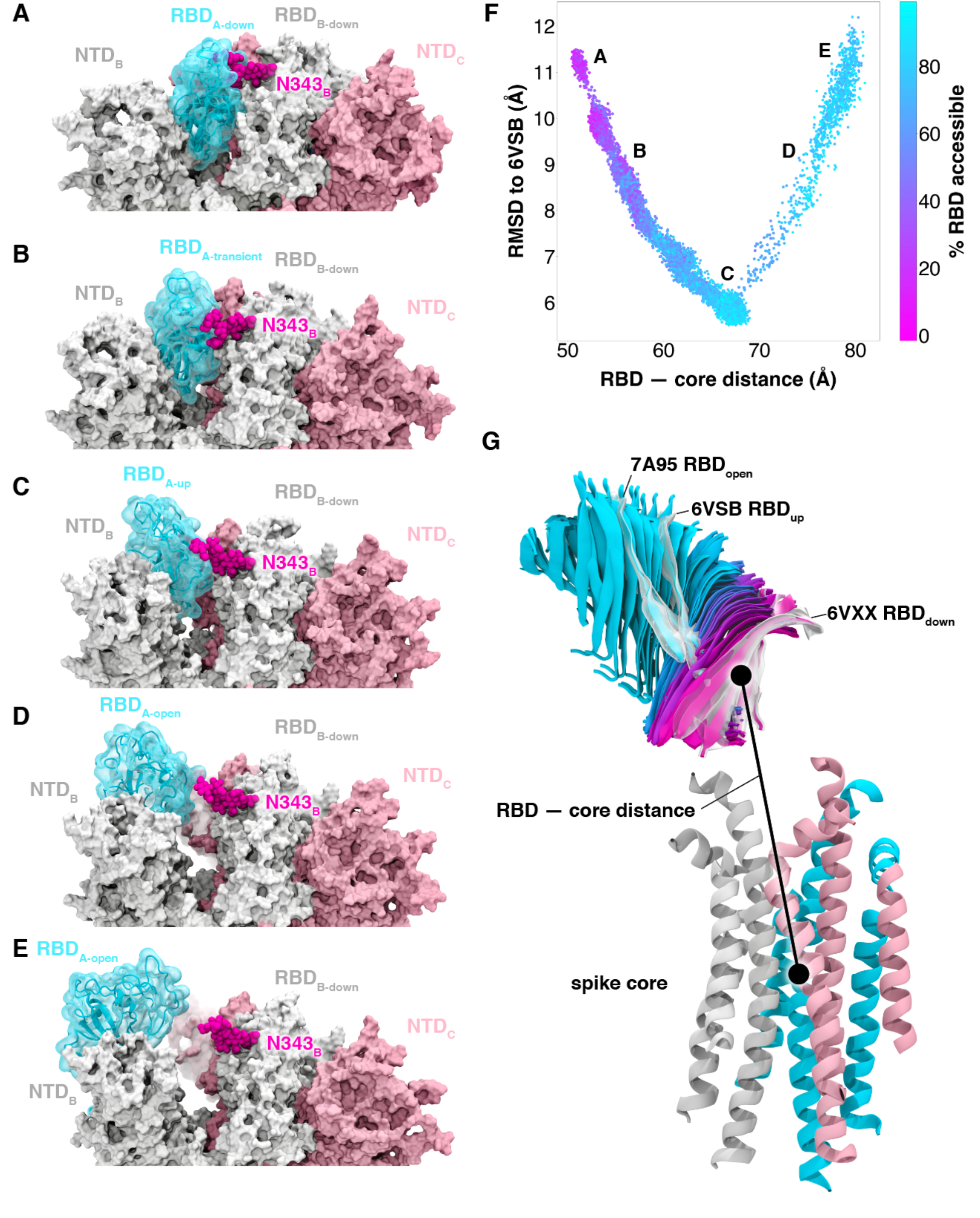
Atomically detailed pathways of spike opening. (A-E) Snapshot configurations along the opening pathway with chain A shown in cyan, chain B in gray, and chain C in pink and the glycan at position N343 is shown in magenta. (F) Scatter plot of data from the 310 continuous pathways with the Cα RMSD of the RBD from the 6VSB “up” state plotted against the RBD — core distance. Data points are colored based on % RBD solvent accessible surface area compared to the RBD “down” state 6VXX. Location of snapshots shown in A-E are labeled. (G) Primary regions of spike defined for tracking progress of the opening transition. The spike core is composed of three central helices per trimer, colored according to chain as in (A-E). The RBD contains a structured pair of antiparallel beta sheets and an overlay of snapshots from a continuous WE simulation are shown colored along a spectrum resembling the palette in (F). Overlayed cryoEM structures are highlighted and labeled including the initial RBD “down” state, 6VXX, the target RBD “up” state and the ACE2 bound “open” state, 7A95.

### Comparison to spike conformations detected by ManifoldEM

To validate our simulated RBD-down to RBD-up pathway, the ManifoldEM framework^22^ was applied using the cryo-EM dataset of PDB 6VSB from McLellan and colleagues.^4^ The ManifoldEM method allows characterization of conformational variations as obtained from a single-particle cryo-EM ensemble of a molecule in thermal equilibrium. Two conformational coordinates (CCs) (i.e., collective motion coordinates) were discovered from this dataset, and observed from several exemplary projection directions (PDs) showing a (1) RBD-down to RBD-up pathway and (2) RBD outward opening pathway (**Supplementary Fig. 4 Supplementary Videos 2 and 3**).

These projections were next aligned to corresponding 2D projections of Coulomb potential maps generated with frames from the WE simulation (**Supplementary Fig. 5, Supplementary Videos 2 and 3**). Overall, there was very good agreement between the ManifoldEM conformational coordinates and WE trajectory, aside from two discrepancies. First, the CC2 observed in the ManifoldEM included concerted opening of all three RBDs, while the WE focused sampling on the opening of a single RBD (**Supplementary Video 2**). Second, the WE trajectory ultimately opens to an RBD — core distance 11 Å greater than the most open conformation in the ManifoldEM. This is likely because the simulations sample the S1 subunit *en route* to the post-fusion conformation, whereas the experimental dataset does not.

### The N343 glycan gates RBD opening

In the “down” state, the RBD of the SARS-CoV-2 spike is shielded by glycans at positions N165, N234, and N343.^24^ While glycan shielding had been investigated for the RBD-down and RBD-up states,^8^ our WE simulations allowed characterization of shielding *during* the opening process, revealing an abrupt decrease in glycan shielding when the RBD transitions from the “down” to the “up” state. The glycans at position N165 and N234 consistently shield the receptor binding motif, while shielding by the N343 glycan decreases with RBD opening. (**Supplementary Fig. 6**). Beyond shielding, a structural role for glycans at positions N165 and N234 has been recently reported, stabilizing the RBD in the “up” conformation through a “load-and-lock” mechanism.^8^

Our WE simulations reveal an even more specific, critical role of a glycan in the opening mechanism of the spike: the N343 glycan acts as a “glycan gate” pushing the RBD from the “down” to the “up” conformation by intercalating between residues F490, Y489, F456, and R457 of the ACE2 binding motif in a “hand-jive” motion (**Fig. 2A-E**, **Fig. 3**, **Supplementary Video 4**). Therefore, the N343 glycan plays an active role in initiating the transition, distinct from the stabilizing roles of glycans N165 and N234. This gating mechanism was initially visualized in several successful pathways of spike opening and then confirmed through analysis of all 310 successful pathways in which the N343 glycan was found to form contacts (within 3.5 Å) with each of the aforementioned residues in every successful pathway (**Supplementary Fig. 7**). The same mechanistic behavior of the N343 glycan was observed in two fully independent WE simulations, suggesting the result is robust despite potentially incomplete sampling that can challenge WE and other enhanced sampling simulation methods.^15^

**Figure 3.**
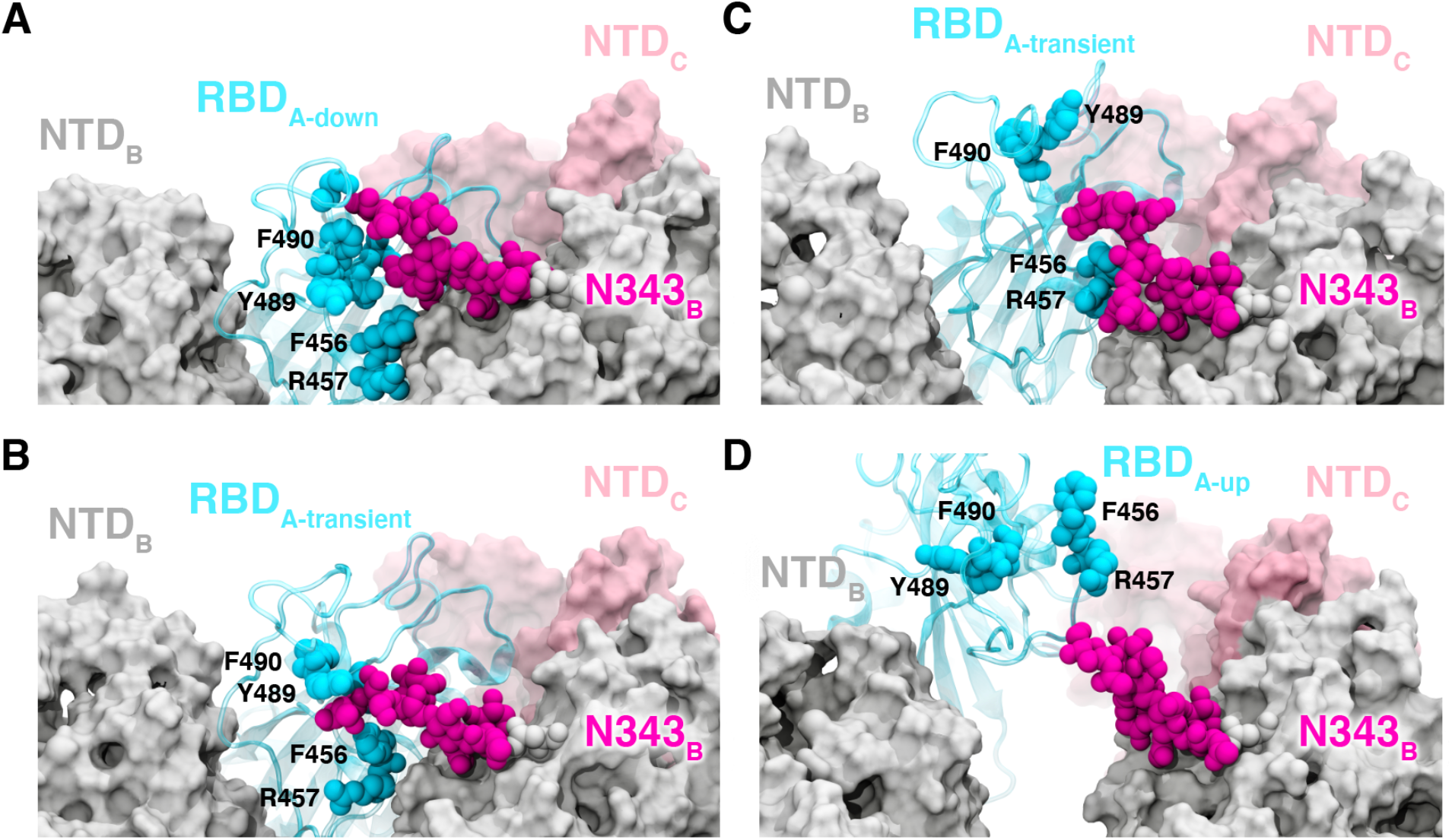
Glycan gating by N343. (A-D) Snapshot configurations along the opening pathway with chain A shown in cyan, chain B in gray, chain C in pink, and the glycan at position N343 is shown in magenta. (A) RBD A in the “down” conformation is shielded by the glycan at position N343 of the adjacent RBD B. (B-D) The N343 glycan intercalates between and underneath the residues F490, Y489, F456, F457 to push the RBD up and open (D).

To test the role of the N343 glycan as a key-gating residue, we performed biolayer interferometry (BLI) experiments. BLI experiments assess the binding level of the spike receptor binding motif (RBM, residues 438 to 508) to ACE2, acting as a proxy for the relative proportion of RBDs in the “up” position for each spike variant. No residues directly involved in the binding were mutated (i.e., at the RBM-ACE2 interface), to ensure controlled detection of the impact of RBD opening in response to mutations. Although previous results have shown reduced binding levels for N165A and N234A variants in the SARS-CoV-2 S-2P protein,^8^ the N343A variant displayed an even greater decrease in ACE2 binding, reducing spike binding level by ~56 % (**Fig. 4, Supplementary Table 1)**. As a negative control, the S383C/D985C variant,^25^ which is expected to be locked by disulfides into the three-RBD-down conformation, showed no association with the ACE2 receptor. These results support the hypothesis that the RBD-up conformation is significantly affected by glycosylation at position N343.

**Figure 4.**
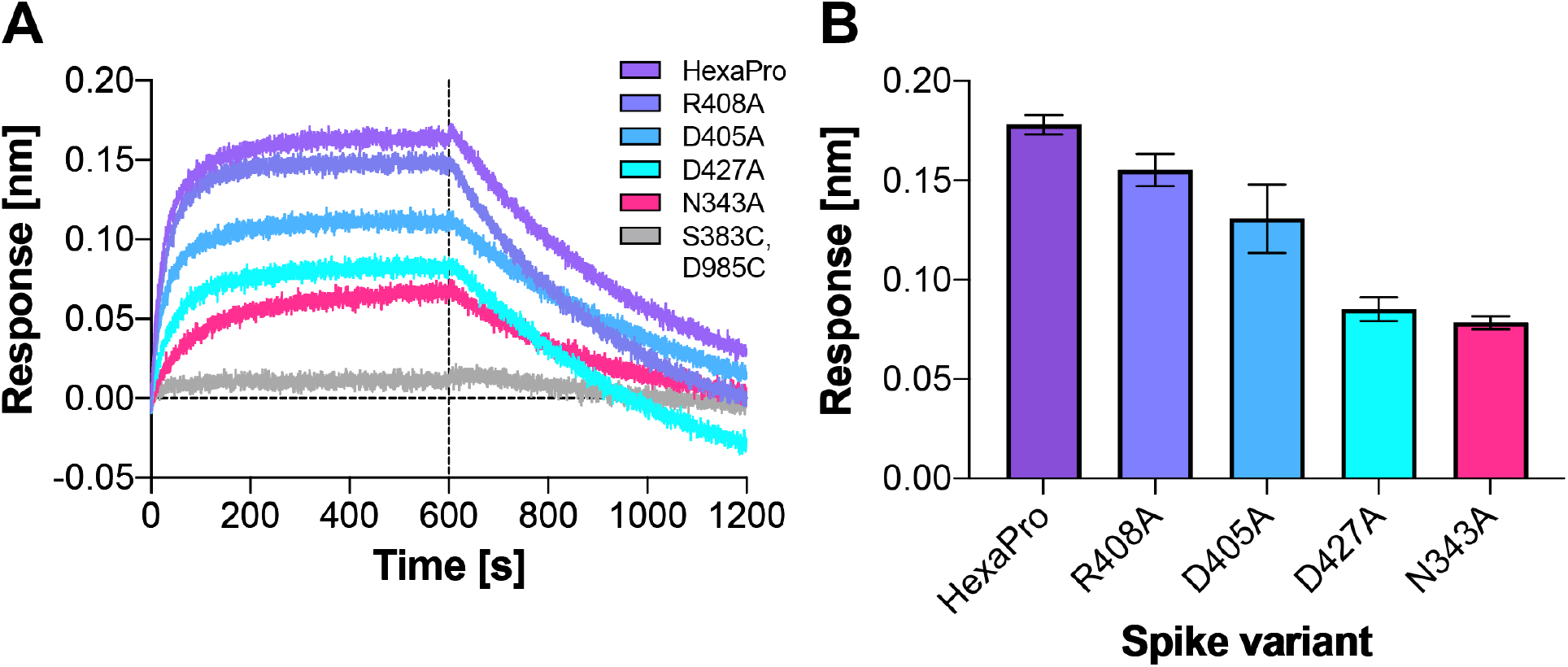
ACE2 binding is reduced by mutation of N343 glycosylation site and key salt bridge residues. (A) Biolayer interferometry sensorgrams of HexaPro spike variants binding to ACE2. For clarity, only the traces from the first replicate are shown. (B) Graph of binding response for BLI data collected in triplicate with error bars representing the standard deviation from the mean.

### Atomic details of the opening mechanism

The RBD-down state features a hydrogen bond between T415 of the RBD (chain A) and K986 of chain C, a salt bridge between R457 of RBDA and D364 of RBDB, and a salt bridge between K462 of RBDA and D198 of NTDC (**Fig. 5A-C,E Supplementary Fig. 8**). The hydrogen bond T415A - K986C spends an average of 12% of the successful pathways to the “up” state before K986C makes a short lived (2% average duration to “up” state) salt bridge with RBDA D427. (**Fig. 5B,E Supplementary Fig. 8**) Next, K986C forms salt bridges with E990C and E748C as the RBDA continues to open. These contacts are formed in all 310 successful pathways (**Supplementary Fig. 8**). Mutation of K986 to proline has been used to stabilize the pre-fusion spike,^23,26^ including in vaccine development,^27^ and these simulations provide molecular context to an additional role of this residue in RBD opening.

**Figure 5.**
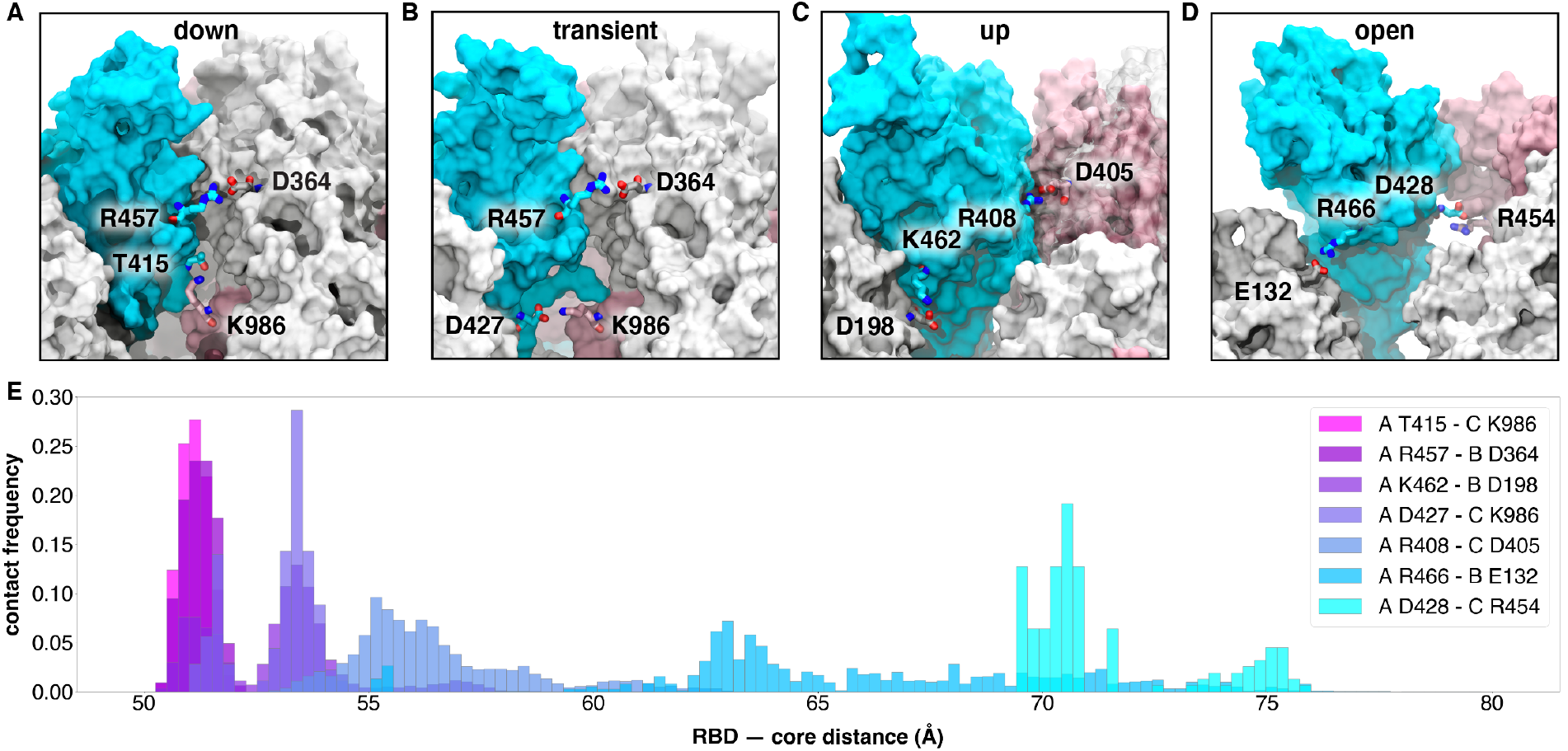
Salt bridges and hydrogen bonds along the opening pathway. (A-D) salt bridge or hydrogen bond contacts made between RBD A, shown in blue, with RBD B, shown in gray, or RBD C shown in pink within the “down”, transient, “up”, and “open” conformations, respectively. (E) Histogram showing the frequency at which residues from (A-D) are within 3.5 Å of each other relative to RBD — core distance. Frequencies are normalized to 1.

Subsequently, at an average of 16% of the way through the successful pathways to the “up” state, the R457A - D364B salt bridge is broken, prompting the RBDA to twist upward, away from RBDB towards RBDC and forming a salt bridge between R408 of RBDA and D405 of RBDC (**Figs. 5C and 5E, Supplementary Fig. 8**). This salt bridge persists for 20% of the successful trajectories to the “up” state and is present in all 310 successful pathways.

A salt bridge between R466 of RBDA and E132 from NTDB is present in 189 out of 204 successful pathways to the “up” state, and all 106 pathways to the “open” state. This contact is most prevalent during the transition between the “up” and “open” state. Finally, the salt bridge between D428 of RBDA and R454 of RBDC is present only in all 106 pathways from the “up” to the “open” state and is the last salt bridge between the RBD and the spike in the “open” state before the S1 subunit begins to peel off (**Fig. 5D and 5E, Supplementary Fig. 8**), at which point the last remaining contact to the RBDA is the glycan at position N165 of NTBB.

Additional BLI experiments of the key identified spike residues R408A, D405A, and D427A corroborate the pathways observed in our simulations. Each of these reduces the binding interactions of the spike with ACE2 by ~13%, ~27%, and ~52%, respectively (**Fig. 4, Supplementary Table 1**). We also note that identified residues D198, N343, D364, D405, R408, T415, D427, D428, R454, R457, R466, E748, K986 and E990 are conserved between SARS-CoV and SARS-CoV-2 spikes, supporting their significance in coordinating the primary spike function of RBD opening. The emerging mutant SARS-CoV-2 strains B.1: D614G; B.1.1.7: H69-V70 deletion, Y144-Y145 deletions, N501Y, A570D, D614G, P681H, T716I, S982A, D1118H; B.1.351: L18F, D80A, D215G, R246I, K417N, E484K, N501Y, D614G, A701V; P1: L18F, T20N, P26S, D138Y, R190S, K417T, E484K, N501Y, D614G, H655Y, T1027I and CAL.20C: L452R, D614G^28^ do not contain mutants in the residues we identified here to facilitate RBD opening. Analysis of neighboring residues and glycans to those mutated in the emerging strains along the opening pathway is detailed in **Supplementary Table 2**, and distances between each residue and glycan to RBDA is summarized in **Supplementary Video 5**.

## Conclusions

We report extensive weighted ensemble molecular dynamics simulations of the glycosylated SARS-CoV-2 spike head characterizing the transition from the “down” to “up” conformation of the RBD. Over 130 microseconds of simulation provide more than 300 independent RBD opening transition pathways. The simulated opening pathways align very well to conformations detected from cryoEM with the ManifoldEM method. Analysis of these pathways from independent WE simulations indicates a clear gating role for the glycan at N343, which lifts and stabilizes the RBD throughout the opening transition. We also characterize an “open” state of the spike RBD, in which the N165 glycan of chain B is the last remaining contact with the RBD *en route* to further opening of S1. Biolayer interferometry experiments of residues identified as key in the opening transitions, including N343, D405, R408, and D427, broadly supported our computational findings. Notably, a 56% decrease in ACE2 binding of the N343A mutant, compared to a 40% decrease in N234A mutant, and 10% decrease in the N165A mutant reported previously^8^ evidenced the key role of N343 in gating and assisting the RBD opening process, highlighting the importance of sampling functional transitions to fully understand mechanistic detail. None of the individual mutations fully abolished ACE2 binding, indicating the virus has evolved a mechanism involving multiple residues to coordinate spike opening. Our work indicates a critical gating role of the N343 glycan in spike opening and provides new insights to mechanisms of viral infection for this important pathogen.

## Acknowledgements

We are grateful for the efforts of the Texas Advanced Computing Center (TACC) Longhorn team and for the compute time made available through a Director’s Discretionary Allocation (made possible by the National Science Foundation award OAC-1818253). We thank Dr. Zied Gaieb for helpful discussions around system construction. We thank Mahidhar Tatineni for help with computing on SDSC Comet, as well as a COVID19 HPC Consortium Award for compute time. We also thank Prof. Carlos Simmerling and his research group (SUNY Stony Brook), and Prof. Adrian Mulholland and his research group (University of Bristol) for helpful discussions related to the spike protein, as well as Prof. Daniel Zuckerman, Dr. Jeremy Copperman, Dr. Matthew Zwier, and Dr. Sinam Saglam for helpful methodological discussions.

## Funding

T.S. is funded by NSF GRFP DGE-1650112. This work was supported by NIH GM132826, NSF RAPID MCB-2032054, an award from the RCSA Research Corp., and a UC San Diego Moores Cancer Center 2020 SARS-COV-2 seed grant to R.E.A.; NIH grant R01-GM31749 to J.A.M.; NIH grant R01-AI127521 to J.S.M; NIH R01 GM115805 and NSF CHE-1807301 to L.T.C; and NIGMS R01 GM29169 and R35 GM139453 to J.F.

## Author contributions

T.S. and S.A. contributed equally to this work. R.E.A. and L.T.C. oversaw the project. T.S. and L.C. prepared the simulation model. T.S. and S.A. performed WE simulations and A.B. provided WESTPA scripts. S.A., A.B., T.S., and L.T.C. carried out WE analysis. T.S., and F.L.K. performed simulation analyses. L.C., T.S., and F.L.K. created figures and movies. J.S.M. designed and oversaw biolayer interferometry experiments. R.S.M. and J.A.G. performed biolayer interferometry experiments and wrote the corresponding parts in the Results and Methods sections. J.F. and A.O. directed the ManifoldEM study, E.S., F.A., S.M. and G.M. performed the ManifoldEM study. E.S. and F.A. analyzed the results and compared them with WE simulations. E.S. and J.F. described the ManifoldEM methods and results. T.S., S.A., L.T.C., and R.E.A. wrote the manuscript with contributions from all authors.

## Competing interests statement

The authors declare no competing financial interests.

## Data availability

Data supporting the findings of this study are included in the Article and its Supplementary Information files. We endorse the community principles around open sharing of COVID19 simulation data.^29^ All simulation input files and data are available at the NSF MolSSI COVID19 Molecular Structure and Therapeutics Hub at https://covid.molssi.org and the Amaro Lab website http://amarolab.ucsd.edu.

## Code availability

This study utilized the standard builds of the simulation software WESTPA 2020.02 (https://github.com/westpa/westpa) and AMBER 18 (https://ambermd.org) according to best practices for running WE simulations^30^ with no special modifications.

## Supplementary information

Sztain_Ahn_Supplementary_Information.pdf

Supplementary_Table2.xlsx

Supplementary_Video1.mp4

Supplementary_Video2.mp4

Supplementary_Video3.mp4

Supplementary_Video4.mp4

Supplementary_Video5.mp4

**Source data**

Source_data_fig2.zip

Source_data_fig4.zip

Source_data_fig5.zip

## Notes

https://amarolab.ucsd.edu/files/covid19/TRAJECTORIES_continuous_spike_opening_WE_chong_and_amarolab.tar.gz

### Competing Interest Statement

The authors have declared no competing interest.

