## Supporting Information for "A glycan gate controls opening of the SARS-CoV-2 spike protein"

**Table of Contents**

1. **Supplementary Methods**

   **1.1 Computational Methods**

   - **1.1.1** Model preparation of the initial “down” state
   - **1.1.2** Weighted ensemble simulations
   - **1.1.3** Analysis of weighted ensemble simulations

   **1.2 ManifoldEM Methods**

   - **1.2.1** Background
   - **1.2.2** Preprocessing
   - **1.2.3** Manifold embedding
   - **1.2.4** Comparison of WE simulations to manifold outputs

   **1.3 Experimental Methods**

2. **Supplementary Figures 1 – 16**
3. **Supplementary Tables 1 – 2**
4. **Supplementary Videos 1 – 5**
5. **Supplementary References**

### 1. Supplementary Methods

#### 1.1 Computational Methods

##### 1.1.1 Model preparation of the initial "down" state

A model of the "down" state of the glycosylated spike structure and CHARMM36 force field parameters[31,32] was obtained from Casalino *et al*,[8] modeled using the cryoEM structure (PDB ID: 6VXX);[5] in this model hydrogen atoms were added using ionization states present in solution at pH 7.4. The stalk and membrane were excluded, and only residues 16-1140 of each trimer were used (**Fig. 1A**). The system was solvated in a cubic box of TIP3P[33] explicit water molecules with at least 10 Å between the protein and box edges and 150 mM NaCl using VMD,[34] yielding a system size of 490,621 atoms. The GPU-accelerated Amber18[35,36,37,38] molecular dynamics (MD) engine was used, which gave a 16-fold speedup in dynamics propagation on a GPU vs. CPU. To enable the use of the Amber18 software package, the Chamber program[39] was used to convert the CHARMM36 force field parameters into an Amber readable format.

To relieve unfavorable interactions, the solvated system was subjected to a two-stage energy minimization followed by a two-stage equilibration. To minimize the energy of the system, the solvent was first minimized for 10,000 steps with harmonic position restraints (force constant of 100 kcal/mol /$Å^2$) applied to the sugars and proteins followed by an unrestrained minimization of the entire system for 100,000 steps. To equilibrate the energy-minimized system, the system was incrementally heated to 300 K over 300 ps in the NVT ensemble followed by a 1-ns equilibration in the NPT ensemble. A production simulation was then carried out in the NPT ensemble for 20 ns on the Triton Shared Computing Cluster at San Diego Supercomputer Center (SDSC). Equilibration and production simulations were carried out with a 2 fs timesteps and SHAKE[40] constraints on bonds to hydrogens. Pressure and temperature were controlled with the Monte Carlo barostat (with 100 fs between attempts to adjust the system volume) and the Langevin thermostat (1 $ps^{-1}$ collision frequency), respectively. Long-range electrostatics were accounted for with the PME method[41] using a 10 Å cutoff for short-range, non-bonded interactions. To provide more extensive sampling of the closed state, we selected a set of 24 equally weighted conformations ("basis states") from the latter 5 ns of the production simulation for a weighted ensemble (WE) simulation; this portion of the simulation exhibited reasonable convergence of the Cα root-mean-squared deviation (RMSD) from the initial, minimized conformation (**Supplemental Fig. 6**).

##### 1.1.2 Weighted ensemble simulations

The weighted ensemble (WE) path sampling strategy orchestrates an ensemble of parallel trajectories with periodic communication to enhance the sampling of pathways for rare events without biasing the dynamics.[15] In particular, a resampling step is applied at fixed time intervals $\tau$ to enrich for promising trajectories that have advanced towards the target state – typically, along a progress coordinate that has been divided into bins. Trajectories are all initially assigned equal statistical weights and rigorously tracked to ensure that all weights sum to one at all times of the simulation, introducing no bias in the dynamics.[12] During the resampling step, trajectories that transition to empty bins are replicated and their corresponding weights split evenly between the resulting child trajectories; trajectories that do not make progress are occasionally terminated with their respective weights merged to other trajectories that will be continued. (**Supplemental Fig. 1**)

WE simulations can be run under non-equilibrium steady state or equilibrium conditions and can therefore provide equilibrium (*e.g.*, state populations) and non-equilibrium observables (e.g., rate constants), respectively. To maintain non-equilibrium steady-state conditions, trajectories that reach the target state are "recycled" by initiating a new trajectory from the initial state with the same trajectory weight; steady-state WE simulations therefore require that the target state be defined in advance of the simulation, but are more efficient in generating successful events than equilibrium WE simulations. On the other hand, equilibrium WE simulations do not require a fixed definition of the target state and therefore enable refinement of the target-state definition at any time during the simulation. Here, we leveraged the advantages of both non-equilibrium steady state and equilibrium WE simulations: steady-state simulations were used to more efficiently generate successful pathways trajectories once the target state could be defined and equilibrium simulations were used to further explore and refine the definition of the target state.

All WE simulations were run using the open-source, highly scalable WESTPA software package[42] (**Supplemental Fig. 7**) with a fixed time interval $\tau$ of 100 ps for resampling and a target number of 8 trajectories/bin. Details of the progress coordinate and bin spacing for each WE simulation are provided below.

*Extensive sampling of the initial "down" state*

To extensively sample the initial "down" state, we ran an equilibrium WE simulation starting from randomly selected conformations from the basis states discussed above. A two-dimensional progress coordinate was used. One dimension consisted of the distance between the centers of mass (COM) of (i) Cα atoms of the entire system **and** all atoms in the four main beta strands of the RBD (residues 375-380, 394-404, 431-438, 508-517; refers to RBD from chain A unless otherwise specified), and (ii) Cα atoms of the entire system **and** all atoms in the structured region of the helical core domain (residues 747-784, 946-967, 986-1034 from each of the three chains). The second dimension consisted of the Cα RMSD of the entire system and all atoms in the four main beta strands of the RBD from the initial model of the "down"-state structure after 1 ns equilibration. Progress coordinates were calculated using CPPTRAJ.[43] This initial WE simulation was run for 8.77 days on 80 P100 GPUs on Comet at the San Diego Supercomputer Center (SDSC) collecting a comprehensive sampling of ~7.5 μs aggregate simulation time. Bin spacing was periodically monitored and adjusted to maximize efficient sampling.

Due to a typo in the CPPTRAJ atom selection *(i.e.,* "and" instead of "of"), the progress coordinate above was not the one we originally intended. Our intention was to use 1) the COM distance between the Cα atoms **of** the four main beta sheets of the RBD and the Cα atoms of the structured region of the helical core domain and 2) the Cα RMSD **of** the four main beta sheets of the RBD from the initial model of the "down"-state structure. As shown in **Figs. 2F and S2**, our WE simulations with this progress coordinate nonetheless capture the large-scale protein transitions that are evident with the intended progress coordinate, but on a more compressed scale.

*Simulations of spike opening*

After extensive sampling of the "down" state, exploratory WE simulations were run to determine effective progress coordinates and binning to capture the opening of the spike protein. Based on these simulations, we found that taking the RMSD from the target "up" state was much more effective than taking the RMSD from the initial "down" state. The target state, with one RBD in the "up" conformation, modeled by Casalino *et al.*[8] using the cryoEM structure (PDB ID: 6VSB),[4] was subject to 1 ns of equilibration using identical methods as described above for the closed structure. The RMSD of the initial state from the target state was calculated as 11.5 Å.

Next, an independent, equilibrium WE simulation was conducted using the two-dimensional progress coordinate described above for sampling the "down" state, but taking the RMSD from the target "up" state instead of the initial "down" state and using the bin spacing determined by the exploratory simulations. The WE simulation was stopped for analysis after 1729 iterations, 19.64 days on 100 NVIDIA V100 GPUs on Longhorn at TACC, collecting an aggregate of ~51.5 µs of sampling and 106 pathways from the "down" to the "open" state. Finally, another WE simulation that was under non-equilibrium steady-state conditions was conducted to maximize sampling of transitions from the "down" to the "up" states. This WE simulation started from iteration 1576 of the previous WE simulation, which was the last iteration before the RBD-COM distance was 9.0 Å or greater, was stopped for analysis after 3000 iterations, 25.03 days later, on 100 NVIDIA V100 GPUs on Longhorn at TACC, collecting an additional ~69.2 µs of sampling and 204 pathways from the "down" to the "open" state. The WESTPA software was shown to scale almost linearly on these 100 NVIDIA V100 GPUs on Longhorn (**Supplemental Fig. 7**), which enabled fast and efficient simulation of the spike.

##### 1.1.3 Analysis of weighted ensemble simulations

*Number of successful pathways*

The successful pathways that reached the "up" state (8.9 Å ≤ RBD-COM distance) or the "open" state (9.9 Å ≤ RBD-COM distance) were obtained by counting all arrivals to that particular state at every WE iteration, which yielded 204 and 106 pathways, respectively. We consider these pathways to be statistically independent pathways. The splitting trees for the 204 and 106 pathways, respectively, can be seen in **Supplemental Figs. 8 and 9**, respectively, which shows trajectory segments shared by the pathways and points of splitting the pathways. The number of pathways is similar to that obtained from calculating the autocorrelation function of arrivals to the “up” and “open” states at a particular WE iteration. For instance, at the end of the WE simulation that sampled the “open” state, there were 1824 trajectories in total and 1193 trajectories that were part of the “open”-state ensemble (defined in later sections as 9.0 Å ≤ RBD-COM distance). Out of the 1193 trajectories that reached the “open”-state ensemble, 133 trajectories were calculated to be statistically independent from calculating the autocorrelation function of the number of arrivals to the “open”-state ensemble[19] (**Supplemental Fig. 10**). The correlation time was calculated to be 16 WE iterations or 1.6 ns so the trajectories that did not share a common segment for 16 iterations from the last point in the trajectory were considered to be statistically independent. By checking these multiple independent pathways that reached the “up” or “open” states, we were able to confirm reproducibility of the identified glycan and residue interactions involved in the particular transition. For calculating the shortest and longest transition times, all successful pathways were taken into account. The first 25% of all successful pathways were disregarded to obtain the most probable transition times, however, since the initial transitions can skew the transition time to be shorter than it is normally (**Supplemental Figs. 11 and 12**).

*State definitions*

Based on our WE simulations, key states were defined as follows. The “down”-state ensemble consisted of structures with RMSD ≥ 11.0 Å and RBD-COM distance ≤ 7.5 Å, ~13 μs aggregate simulation time. Note that the entire progress coordinate array had to satisfy the criteria to be counted as part of the ensemble. The “up”-state ensemble was defined as 8.5 Å ≤ RBD-COM distance < 9.0 Å, ~ 6.5 μs aggregate simulation time. The “open”-state ensemble was defined as having an RBD-COM distance ≥ 9.0 Å, ~ 4.9 μs aggregate simulation time.

*Trajectory analysis*

Trajectories were visualized using VMD.[34] Glycans, salt bridge, and hydrogen bonding interactions involved in the “down” to “up” and “open” transition were first visually identified. Next, distances between the identified residues were calculated using cpptraj[43] for all 310 successful pathways, and plotted with matplotlib.[44] To obtain the percentage of the most probable transition time that had a certain salt bridge, the distance between the atoms/residues of the salt bridge was measured, and the total time in which the distance was less than 3.5 Å was calculated. The total time for each pathway was calculated and averaged to obtain the final percentage. To obtain the number of successful pathways that had a certain quantity, *e.g.,* salt bridge, glycan-residue contact, the pathway was counted if the distance was less than 3.5 Å in at least one of the conformations, sampling conformations every 100 ps. Contact maps calculating the distance between the RBD (from chain A) and all other residues and glycans were generated using MDAnalysis[45,46] (**Supplemental Video 5**). Structures for figures and movies were generated using VMD, including NanoShaper[47] surface representation.

Solvent accessible surface area (SASA) was calculated using a protocol presented in Casalino *et al.*[8] involving the *measure sasa* command within VMD and a solvent probe radius of 1.4 Å. The surface area of the Receptor Binding Motif (RBM, residues 438-508 in chain A) that was shielded by glycans was calculated by taking the difference between the SASA of the "naked" spike (without glycans) and the SASA of the glycosylated spike (with glycans). Individual contributions to shielding of the RBM by glycans at positions N165-B, N234-B, N343-B were also calculated by considering only the respective glycans in the SASA calculation of the glycosylated spike.

*Analysis of residues mutated in emerging SARS-CoV-2 strains*

To date, the following SARS-CoV-2 variants have been identified (with mutations to spike noted in parentheticals): B.1 (D614G), B.1.1.7 (H69-V70 deletion, Y144-Y145 deletions, N501Y, A570D, D614G, P681H, T716I, S982A, D1118H), B.1.351: (L18F, D80A, D215G, R246I, K417N, E484K, N501Y, D614G, A701V), P1 (L18F, T20N, P26S, D138Y, R190S, K417T, E484K, N501Y ,D614G, H655Y, T1027I) and CAL.20C (L452R, D614G).[28] To examine potential implications of these mutations on Spike opening mechanics, we have monitored the neighboring residues of key WT residues as a function of the opening mechanism. MDAnalysis[45,46] was used to identify residues whose center of mass was within 10 Å of the center of mass of the key residue of interest. For each contact, the fraction of conformations in the "down", "up", and "open" ensembles containing the contact is provided. Contacts were only considered if they exist within > 5% of all conformations and if the contacting pairs were separated by more than three peptide bonds in one-dimensional sequence.

#### 1.2 ManifoldEM method

##### 1.2.1 Background

The set of algorithms now under the name ManifoldEM[48] employ a three-step procedure[22] to characterize conformational variations in a dataset from single-particle cryo-EM of a molecule in thermal equilibrium. In the first step, which can be performed on any of the existing cryo-EM platforms, data are classified by orientation, and prepared as aligned image stacks. In the second step, for each projection direction (PD) data falling into the angular aperture are analyzed as a manifold and represented in a low-dimensional space spanned by what is now termed "conformational coordinates,"[48] equivalent to collective motion coordinates. In the third step, the manifold representations resulting from the second step, one for each projection direction, are reconciled and combined across the angular sphere to obtain a consolidated representation. From this an energy landscape can be obtained, enabling a functional analysis of the molecule,[48] and 3D volumes can be captured along inferred trajectories.

##### 1.2.2 Preprocessing

The initial image-stack we received from McLellan and colleagues corresponding to PDB ID: 6VSB[4] contained 631,920 snapshots. This initial image stack was pruned by approximately 10% (from 631,920 to 578,588 particles) to remove artifacts. Additional 3D Auto-Refinement via RELION[49] was performed to realign all images. Next RELION 2D Classification was used to remove an additional 1% of particles, leaving the final count of 574,324. The consensus refinement in RELION displayed a Fourier Shell Correlation ($FSC_{0.143}$) of 4.3 Å. In parallel, this stack was separately refined using CryoSPARC[50] non-uniform refinement with a GSFSC resolution of 3.5 Å.

These two refinements were next compared within the preliminary steps of ManifoldEM. Although both reconstructions appeared fine, we found upon closer examination that the RELION refinement encountered a problem of preferred orientations, where thousands of particles had been clumped within nearly the same local area (*i.e.*, nearly identical Euler coordinates) of the 2-sphere. In contrast, the CryoSPARC non-uniform refinement produced much more uniformly-distributed angular assignments, albeit with a lower average occupancy per PD. 2D conformational coordinate movies obtained in ManifoldEM from the CryoSPARC alignment proved superior to those using RELION. While the CryoSPARC alignment was chosen for all subsequent steps in ManifoldEM, the RELION protocol was not altogether without its own merit. We additionally ran RELION focused 3D Classification using the angular alignment from CryoSPARC with a mask around the RBDs. We obtained classes with different configurations of the RBD, including one class in the RBD-"down" conformation (**Supplemental Fig. 4)**. The original study,[4] in contrast, found no such particles - nor did other labs to which the data were sent for further analysis. Importantly, the discovery of these missing particles explains the presence of RBD-"down" volumes constructed along the 3DVA[51] "reaction coordinate" discovered in that study.[4]

##### 1.2.3 Manifold embedding

We next set up a more thorough ManifoldEM analysis using the cryoSPARC alignment. First, a number of initial inputs are required for the ManifoldEM pipeline to tessellate the orientational 2-sphere into a finite number of PDs. These are (1) Pixel size: 1.047 Å; (2) Resolution: 3.5 Å; (3) Object diameter: 335 Å (taken as the maximum width of the average volume); and (4) Aperture index: {1-5}. The aperture index is a flexible parameter that controls the angular width of each PD, such that a larger aperture index corresponds to more images assigned to each PD from a larger region of angular space. After experimenting with several aperture indices and evaluating the corresponding PD statistics and 2D movie qualities, we chose aperture index 5 for all future computations. This measure provided us with 1678 PDs thoroughly spread out in angular space, with a handful of regions with heightened PD-occupancy. When displayed as a histogram, the occupancy of PDs exhibited a chi-squared distribution, with the majority of PDs housing around 230 images and a rightward tail reaching approximately 800 images in the most highly-occupied PD.

Following the ManifoldEM framework, 1678 manifolds were constructed from the images in each corresponding PD via the Diffusion Maps[52] framework. Following Dashti et al.,[22] Nonlinear Laplacian Spectral Analysis (NLSA)[53] was then performed on the eigenvectors of these high-dimensional manifolds to extract a set of possible reaction coordinates from each. In sum, these steps were programmed to produce eight 2D movies per PD, with each 2D movie corresponding to one of the PD-manifold's eigenvectors.

*Conformational analysis*

Upon completion, our task was to next classify the type of motions seen in each 2D movie per PD, noting that not all 2D movies extracted must correspond to valid conformational information; this is especially true of those obtained with smaller singular values. Our approach was to initiate a search to detect all PDs housing 2D movies with above-average visual appearance. In this search, many PD-manifolds were found to have extremely noisy or otherwise insensible information. This was a predictable scenario given the known deficiencies in the dataset[4] (*i.e.*, orientational bias leading to low occupancies in many PDs), and beyond remediation by ManifoldEM. As a result, only a subset of PDs where the images therein met the prerequisites for the manifold embedding approach could be analyzed. Of these above-threshold PDs, we found 216 PDs of the 1678 PDs (13%) with above average quality and 73 high-quality PDs (4%), as judged by visual inspection relative to the whole. Thus, overall, a relatively small percentage of the data as partitioned into these PDs met the prerequisite conditions for displaying the highest-quality conformational variation signals.

We next organized all above-average PDs into 22 well-spaced groups on the 2-sphere, and selected several of the best PDs from each angular region. Detailed analysis was performed on the 64 PDs chosen, including classification of conformational motion type in each of the eight 2D NLSA movies per PD. As shown in **Supplementary Videos 2 and 3**, we predominantly observed two conformational motions: (1) RBD-"down" to RBD-"up"; and (2) trimer-claw close to open, which we call conformational coordinate 1 (CC1) and conformational coordinate 2 (CC2), respectively. However, PDs where a clear distinction existed between CC1 and CC2 were rare. Specifically, CC1 alone could only be clearly established in 31 of 64 PDs (48%); while both CC1 and CC2 were found occupying separate 2D movies in only 6 of 64 PDs (9%). In the remaining PDs, these conformational motions were not cleanly separated but were present in hybrid form.

This discrepancy arises from the nature of our analysis, where we define Euclidean distances between images that are 2D projections of the molecule. As a result, from a given viewing direction, a 3D motion projected onto 2D will appear more or less pronounced than it does in some other, depending on the type of motion and PD. For example, we found that the CC2 trimer claw motion was most pronounced only when observed from the "top-down view", the PD aligned with the axis of the protein's central alpha helices (PD 112).

*Transformation of structures along WE trajectory*

We next aimed to compare the conformational coordinates discovered by ManifoldEM from experimental cryo-EM ensembles with the WE motions observed in the spike-opening trajectory detailed in the main text. To this end, we converted the PDB files from the WE into a collection of 2D projections. We first selected 20 frames from the WE trajectory spanning conformations from the RBD-"down" to the RBD-"up" state. We next imported these files into Chimera[54] along with a coarse 3D map obtained from ManifoldEM to be used for alignment reference. In order to place both frameworks in the same coordinate system for subsequent analysis, we translated and rotated the PDB files to coincide with the ManifoldEM map, using the Chimera fitmap command. Each PDB was then saved in Chimera. Next, these fitted PDBs were re-centered using Phenix[55] pdbtools and converted into MRC-formatted Coulomb potential maps via EMAN2[56] e2pdb2mrc. For this last step, a resolution of 5 Å was chosen based on visual assessment of the EMAN2 outputs relative to those from ManifoldEM. Projections of these 20 MRCs were then taken using the standard projection operator in e2project3d with C1 symmetry in EMAN2. Importantly, the Euler coordinates for these projections were supplied by those representing the 64 ManifoldEM anchors (after correcting for a coordinate transformation from ManifoldEM to ZXZ' convention). Finally, these projections were combined into sequences for each PD to form 64 20-frame 2D movies of the WE trajectory.

##### 1.2.4 Comparison of WE simulations to ManifoldEM outputs

As shown in **Supplementary Videos 2 and 3**, and described in detail within our main text, a striking visual resemblance emerged between conformational motions obtained by WE simulation and experiment. For heightened visual aid, 2D movies from the WE simulation and the ManifoldEM corresponding to the same PD (and RC therein) were next overlaid to directly highlight similarities and differences. For this procedure, we first layered the ManifoldEM movie over a homogenous red backdrop and applied a Linear Dodge blend mode, with a similar effect applied on the WE movie over a blue backdrop (see **Supplemental Fig. 4** for the results of these operations). We next multiplied the ManifoldEM composite image and the WE composite image together. As an outcome of this multiplication, pixels that are white (signal) in both movies retain their whiteness in the composite. In this way, whiteness in the composite movie becomes a qualitative measure of similarity between conforming domains, while non-white regions emphasize differences.

Finally, this overlaying approach was used to estimate the total extent of the RBD motion as expressed in the ManifoldEM and WE frameworks. For this comparison, CC1 from a side view (PD 1386) was chosen based on its highly prominent view of RBD-“up” to RBD-“down” motion. Next, the ManifoldEM movie was time-remapped to align it optimally in time with the motions observed in the corresponding WE movie (**Supplementary Video 2**). Using the multiplication-composite as a guide, it was determined that the ManifoldEM RBD domain reaches its full extent in the “up” position at the 14$^{th}$ frame out of the 20 frames from the WE trajectory, before the WE trajectory moves onward to a more fully open state. With this knowledge, the total difference in conformational extents was estimated at 11 Å as calculated via RBD — core distance.

#### 1.3 Experimental Methods

*Protein Expression and Purification*

Substitutions N343A, D405A, R408A, and D427A were cloned into the HexaPro SARS-CoV-2 spike background.[23] A spike variant with all RBDs locked in the “down” position through the introduction of a disulfide bond was similarly produced through cysteine substitutions at residues S383C and D985C in the HexaPro protein.[25] All variants were expressed through polyethyleneimine-induced transient transfection of FreeStyle 293-F cells (Thermo Fisher). After 4 days, cell supernatant was clarified by centrifugation, passed through a 0.22 µm filter, and purified over StrepTactin resin (IBA). Variants were further purified by size-exclusion chromatography on a Superose 6 10/300 column (GE Healthcare) in a buffer consisting of 2 mM Tris pH 8.0, 200 mM NaCl and 0.02% $NaN_3$. Soluble ACE2 was produced and purified as previously described.[8]

*Biolayer Interferometry*

Anti-foldon IgG was immobilized to an anti-human Fc (AHC) Octet biosensor (FortéBio). Tips were then submerged into the specified HexaPro variants before being subsequently dipped into 200 nM ACE2 to observe variant association, followed by dissociation in buffer consisting of 20 mM Tris pH 7.5, 150 mM NaCl, 1 mg/mL bovine serum albumin, and 0.01% Tween-20. The relative proportion of RBD in an accessible state was quantified based on the binding level as previously described.[8] The S383C, D985C variant was used as a negative control. Data were collected in triplicate and replicate sensorgrams are shown in **Supplemental Fig. 16.**

#### 2. Supplementary Figures

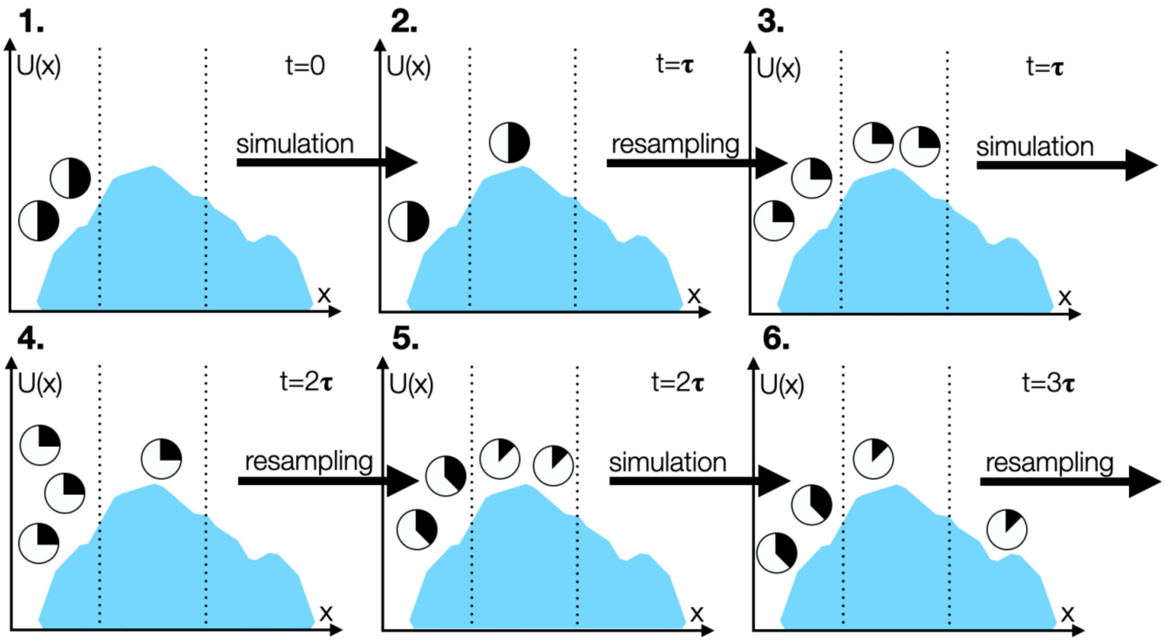

**Supplemental Fig. 1** Schematic of the weighted ensemble (WE) strategy. The WE strategy is illustrated for a three-state system with a one-dimensional progress coordinate x that is divided into bins. U(x) represents the potential of the system dependent on x, which can be seen from the curve of the shaded region. 1. WE initiates two equally weighted trajectories (represented as circles) from the first bin, each with a statistical weight of 0.5 (represented as filled parts of the circles), for a fixed time interval **τ**. 2. Resampling is then performed, replicating or terminating trajectories to maintain a target number of two trajectories in each bin (*e.g.,* in the first and second bins, splitting the weight among the two child trajectories with a weight of 0.25 for each trajectory). 3. Trajectories are run for another fixed time interval **τ**. 4. After running, resampling is performed (*e.g.,* in the first bin, terminating two of the three trajectories and in the second bin, replicating the one trajectory to yield two trajectories). 5. The system ends up with two trajectories in each of the visited bins. 6. One of the trajectories ends up in the third bin. Rounds of simulation and resampling are performed until a desired number of continuous pathways into the target state are generated.

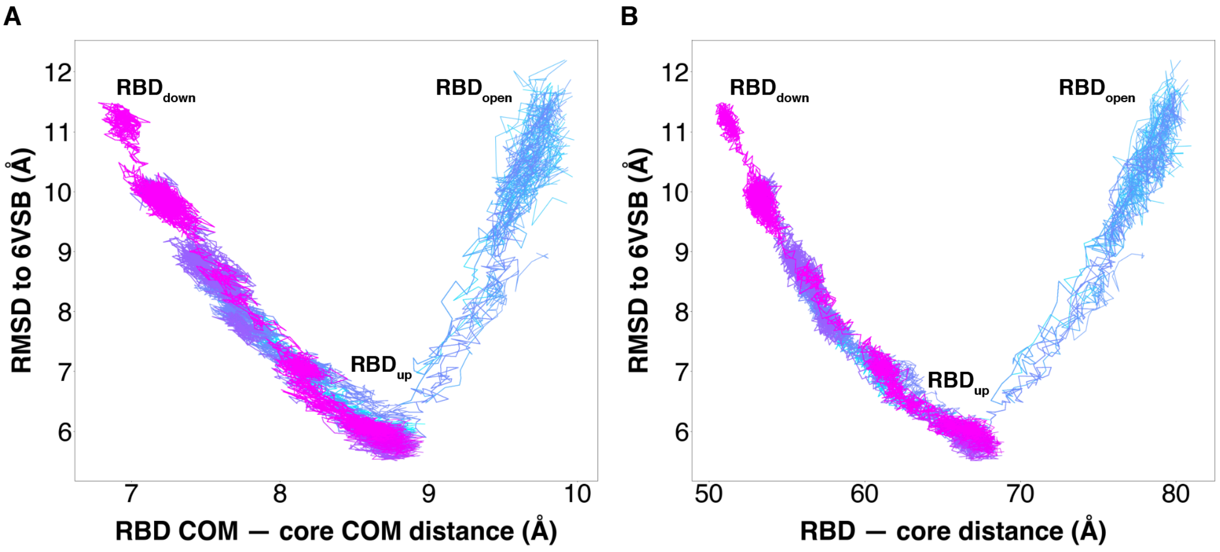

**Supplemental Fig. 2** Successful pathways of spike opening for the (A) actual and (B) intended progress coordinate. Overlay of 310 successful pathways including 204 pathways of the RBD transitioning from the “down” state to the “up” state (magenta-purple) and 106 pathways from the “down” to the “open” states (purple to cyan). Continuous trajectories plotted with the Cα RMSD of the RBD to the 6VSB “up” state versus the RBD — core distance.

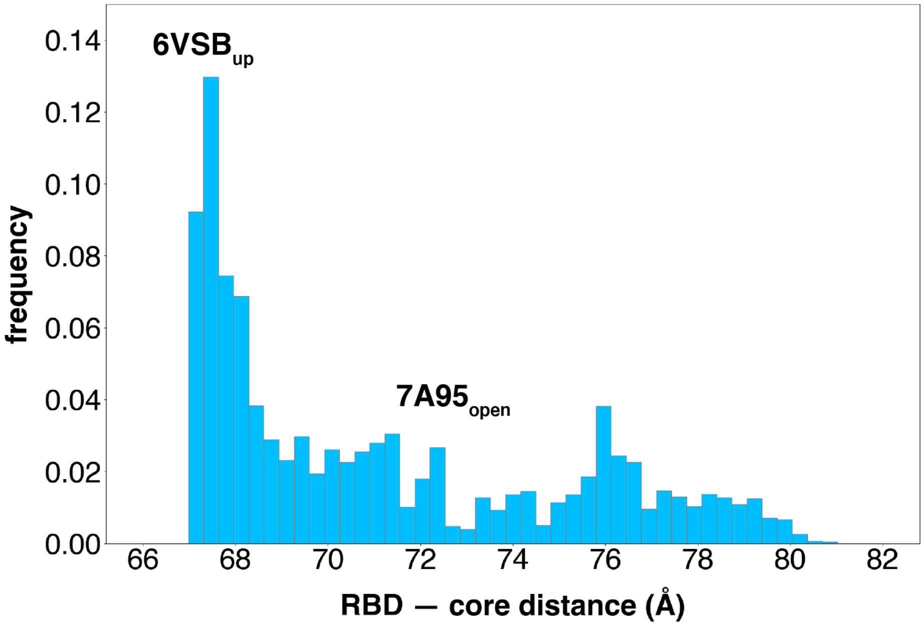

**Supplemental Fig. 3** Diversity of the simulated RBD "open" state ensemble. Probability distribution of RBD — core distances greater than the RBD "up" conformation defined by PDB 6VSB (67.2 Å). The ACE2-bound structure from PDB 7A95 distance is 72.1 Å.

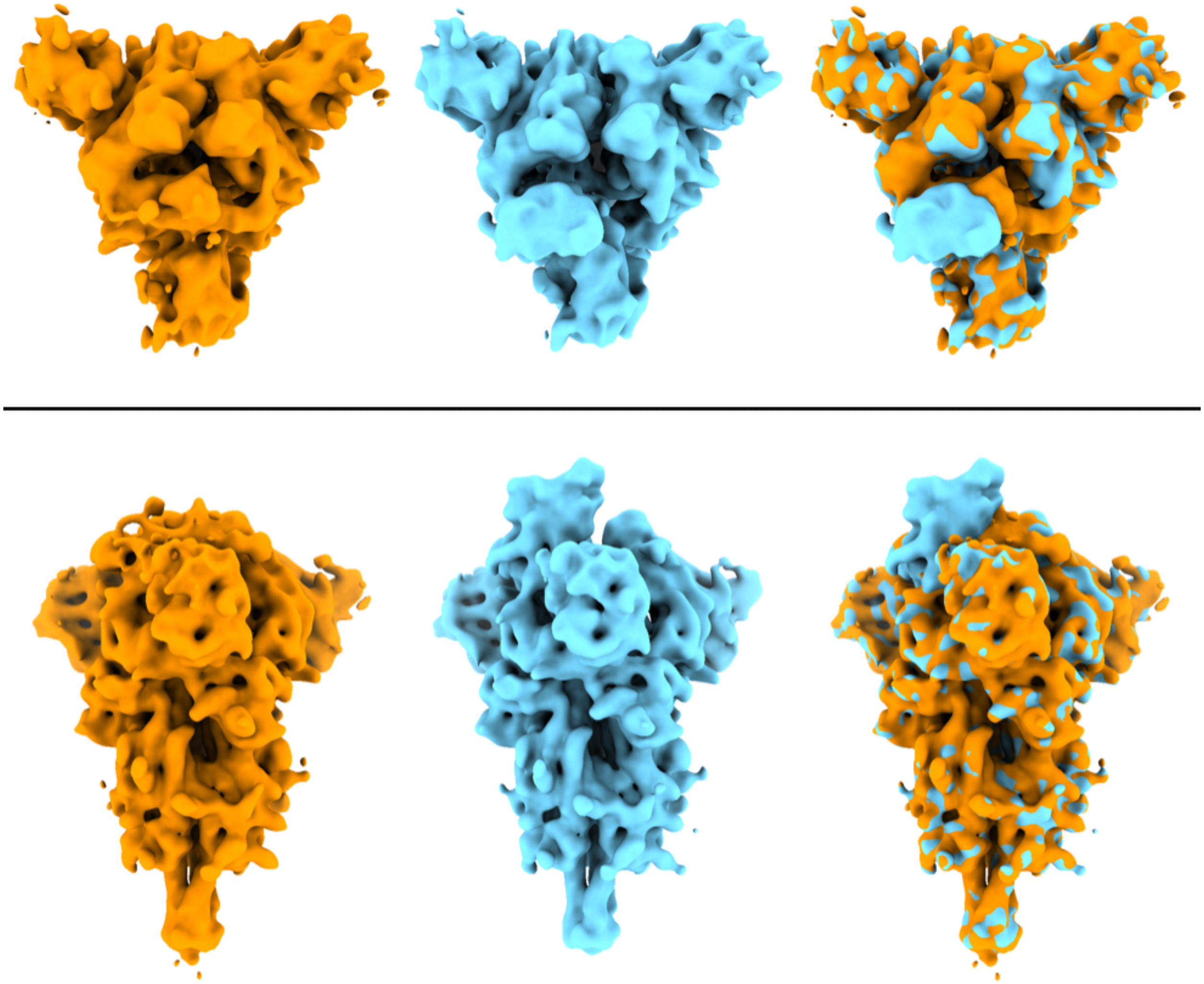

**Supplemental Fig. 4** Comparison of two classes from the focused 3D classification in RELION with top and side views of the reconstructed classes. EM density maps are low pass filtered to 8 Å for display purposes. The class with the RBD "down" conformation is displayed with orange on the left, the class with the RBD "up" is displayed with cyan in the center, and the superposition of both maps is shown on the right side to highlight their differences.

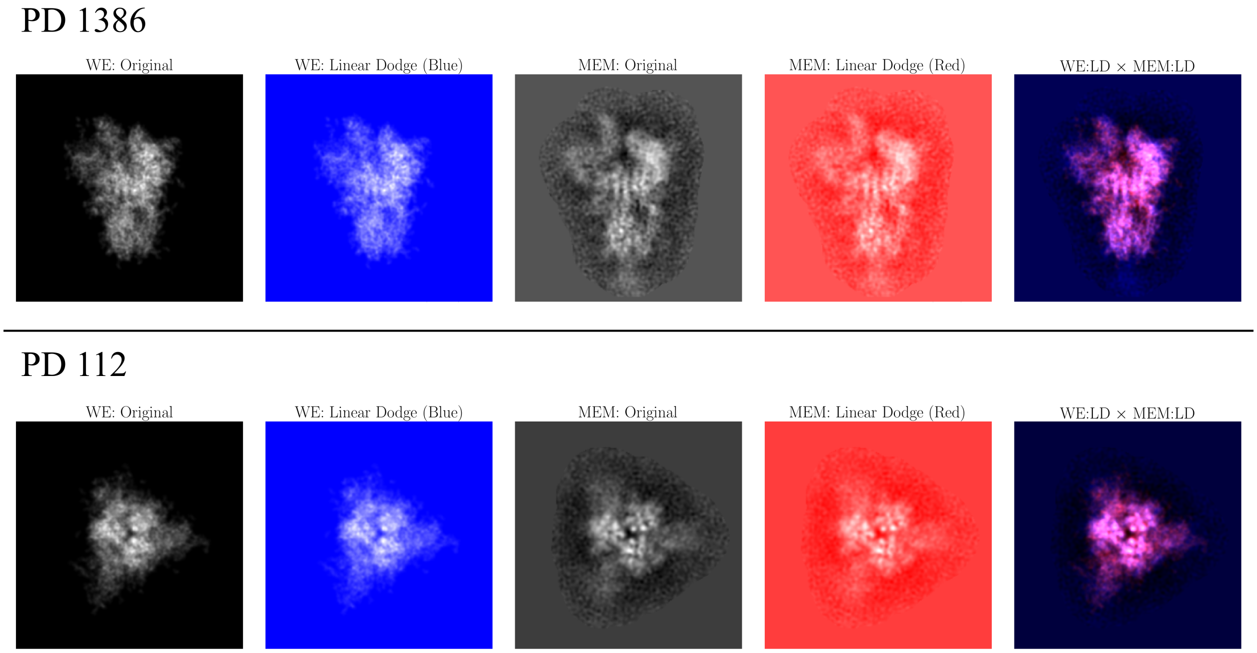

**Supplemental Fig. 5** Comparison of a frame from the WE and ManifoldEM (MEM) trajectory as seen from a side view (PD 1386) and top-down view (PD 112). For this comparison, image compositing techniques are applied on the outputs of each method as shown in the columns, including Linear Dodge and Multiply. As an example of its utility, after performing this operation on RC2 from a top-down view (PD 112), it can be seen that a collection of white pixels emerged in the composite movie (bottom-right entry), which strongly emphasize the similarities in positions of RBD and spike core helices between frameworks.

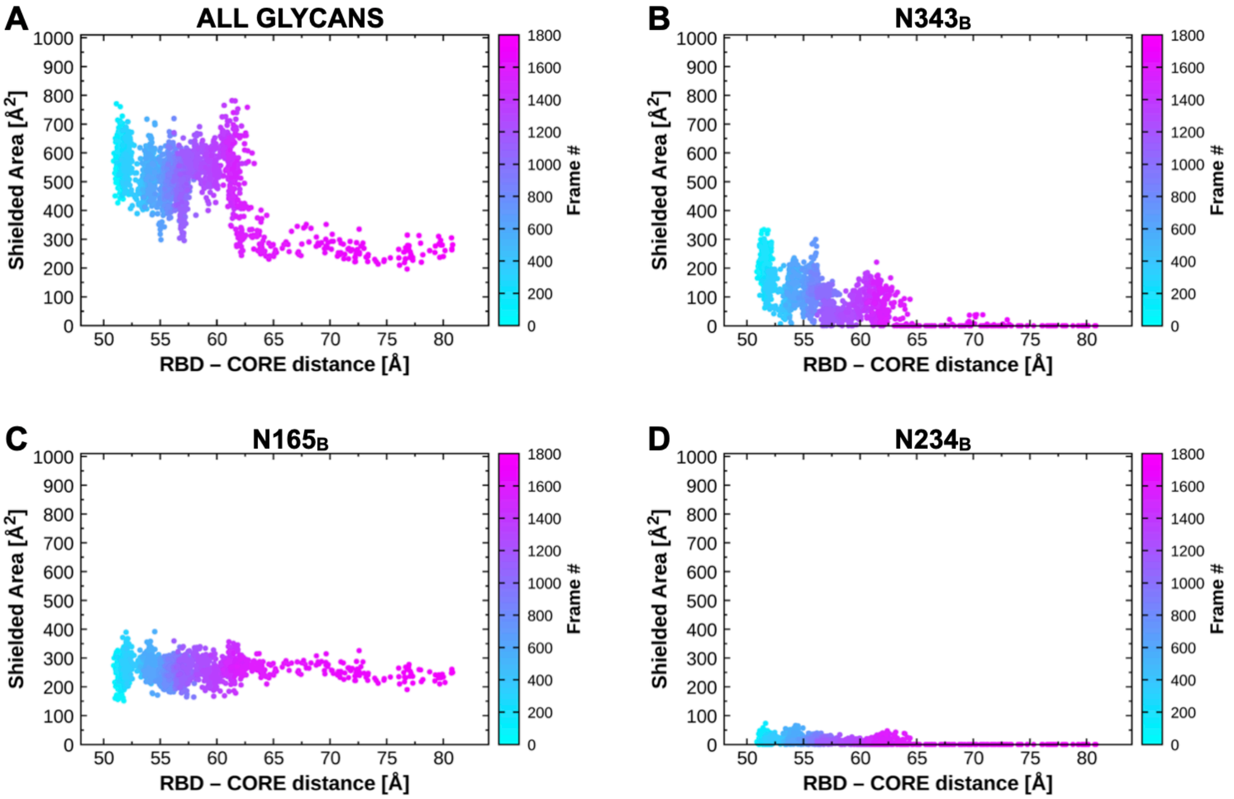

**Supplemental Fig. 6** Contribution of glycans shielding receptor binding motif along RBD opening pathway. Shielded area represents the difference between the solvent accessible surface area of the receptor binding motif in the presence and absence of (A) all three glycans, (B) N343, (C) N165, or (D) N234.

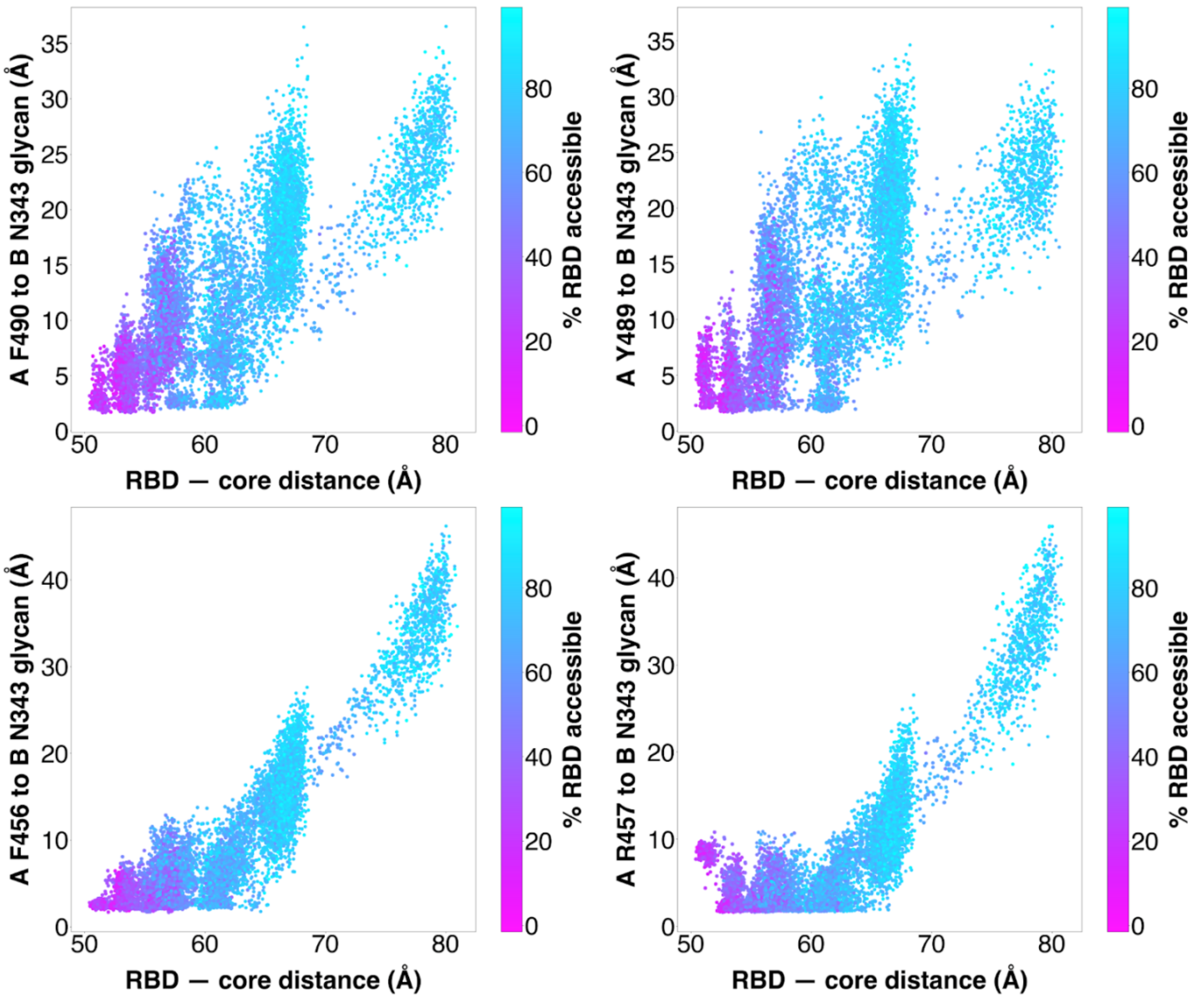

**Supplemental Fig. 7** Distance between N343 glycan and RBD residues. Scatter plot of data from the 310 continuous pathways with the minimum distance between the N343 glycan and RBD A residues F490, Y489, F456, or R457 plotted against RBD — core distance. Data points are colored based on % RBD solvent accessible surface area compared to the RBD "down" state 6VXX.

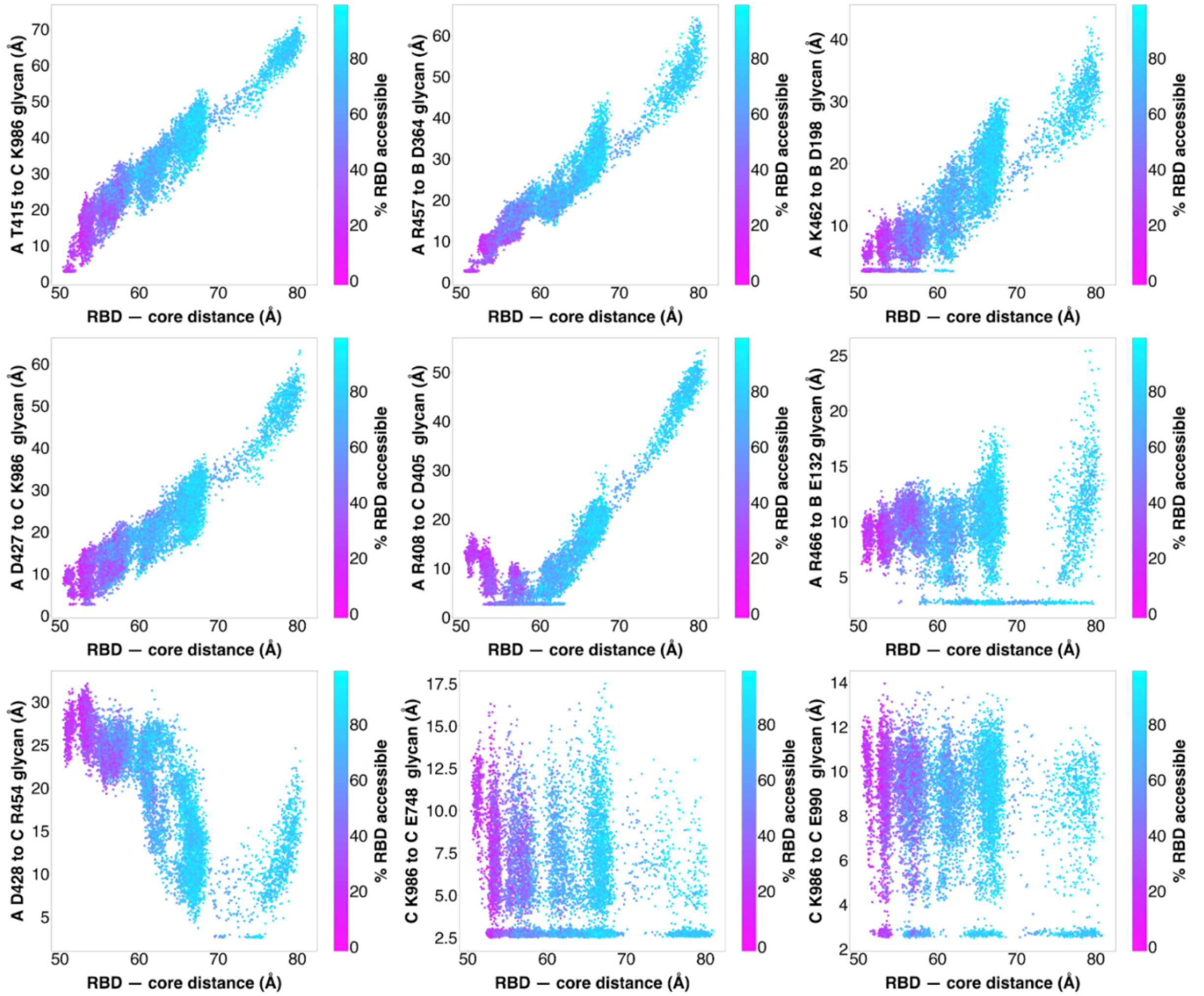

**Supplemental Fig. 8** Distance between salt-bridge and hydrogen bonding residues along the spike opening pathway. Scatter plot of data from the 310 continuous pathways with the minimum distance between the residues shown in **Figure 4** plotted against RBD-core distance. Data points are colored based on % RBD solvent accessible surface area compared to the RBD "down" state 6VXX.

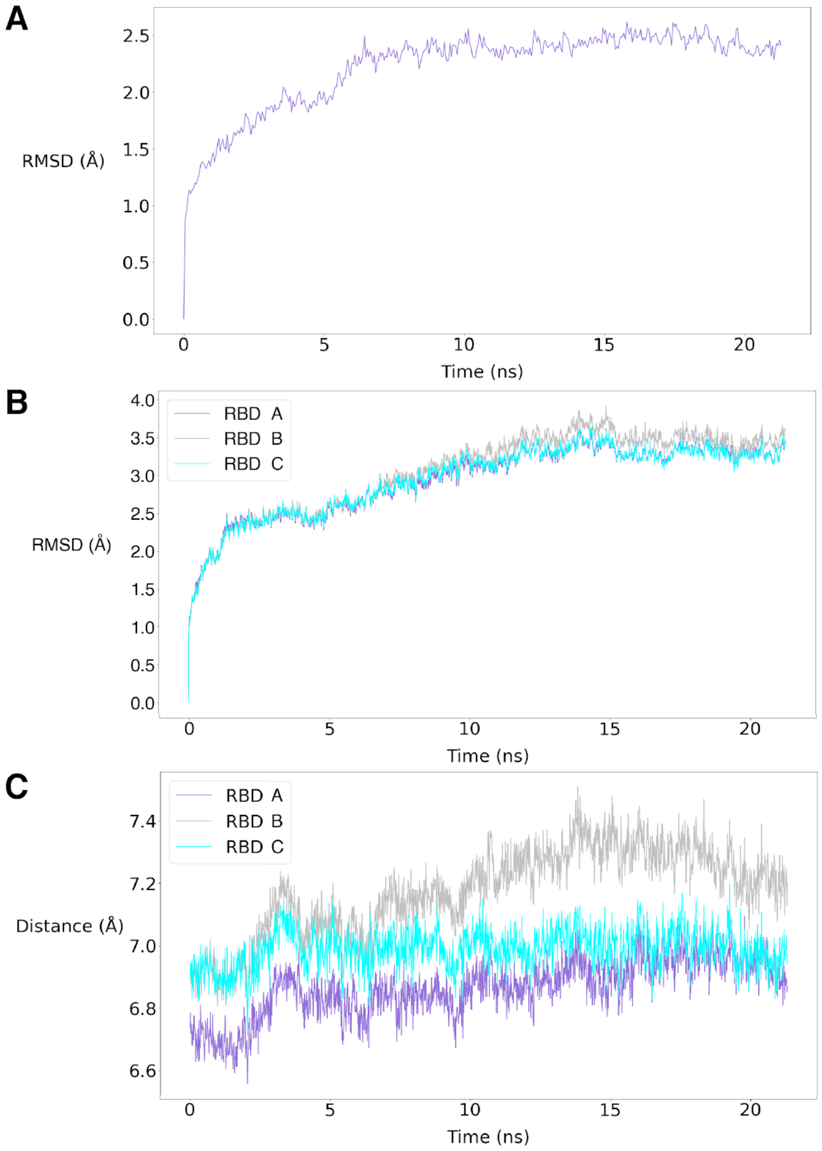

**Supplemental Fig. 9** Initial equilibration of a "down"-state structure using a standard MD simulation. Time evolution of (A) Cα RMSD of protein residues , (B) Cα RMSD of structured region of RBD after alignment of core domain to the initial structure and (C) Distance between centers of mass of the RBD and core domain.

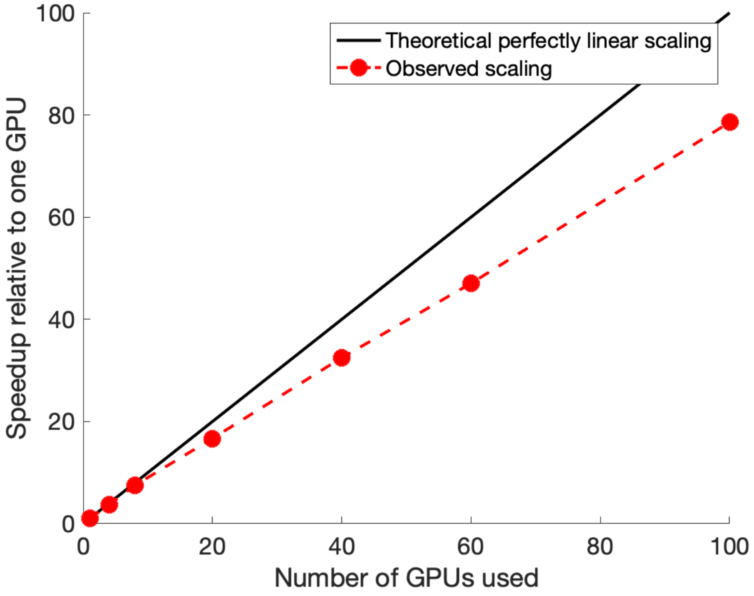

**Supplemental Fig. 10** Scaling of the WESTPA software using NVIDIA V100 GPUs on the TACC Longhorn supercomputer vs. theoretical perfectly linear scaling.

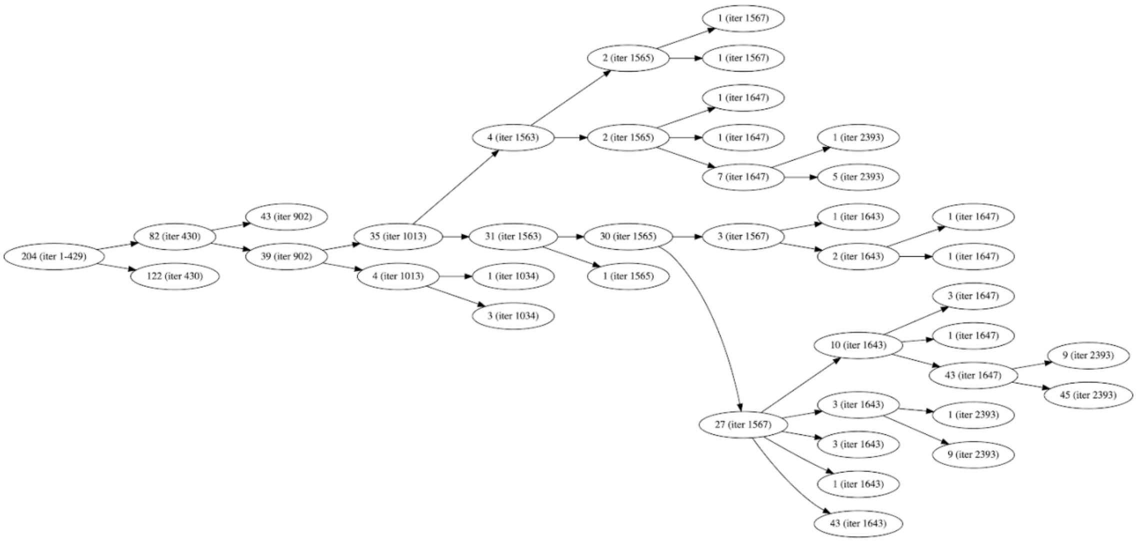

**Supplemental Fig. 11** Trajectory splitting tree of the 204 independent pathways that reached the "up" state. The number of each node indicates the number of pathways at the given WE iteration in parentheses. All trajectories shared the same parents until iteration 429, with the first splitting of trajectories occurring at iteration 430. Subsequent splitting occurred at later iterations. Note that the sum of the child pathways does not necessarily match up with the parent's number of pathways due to splitting and merging with other trajectories (not shown).

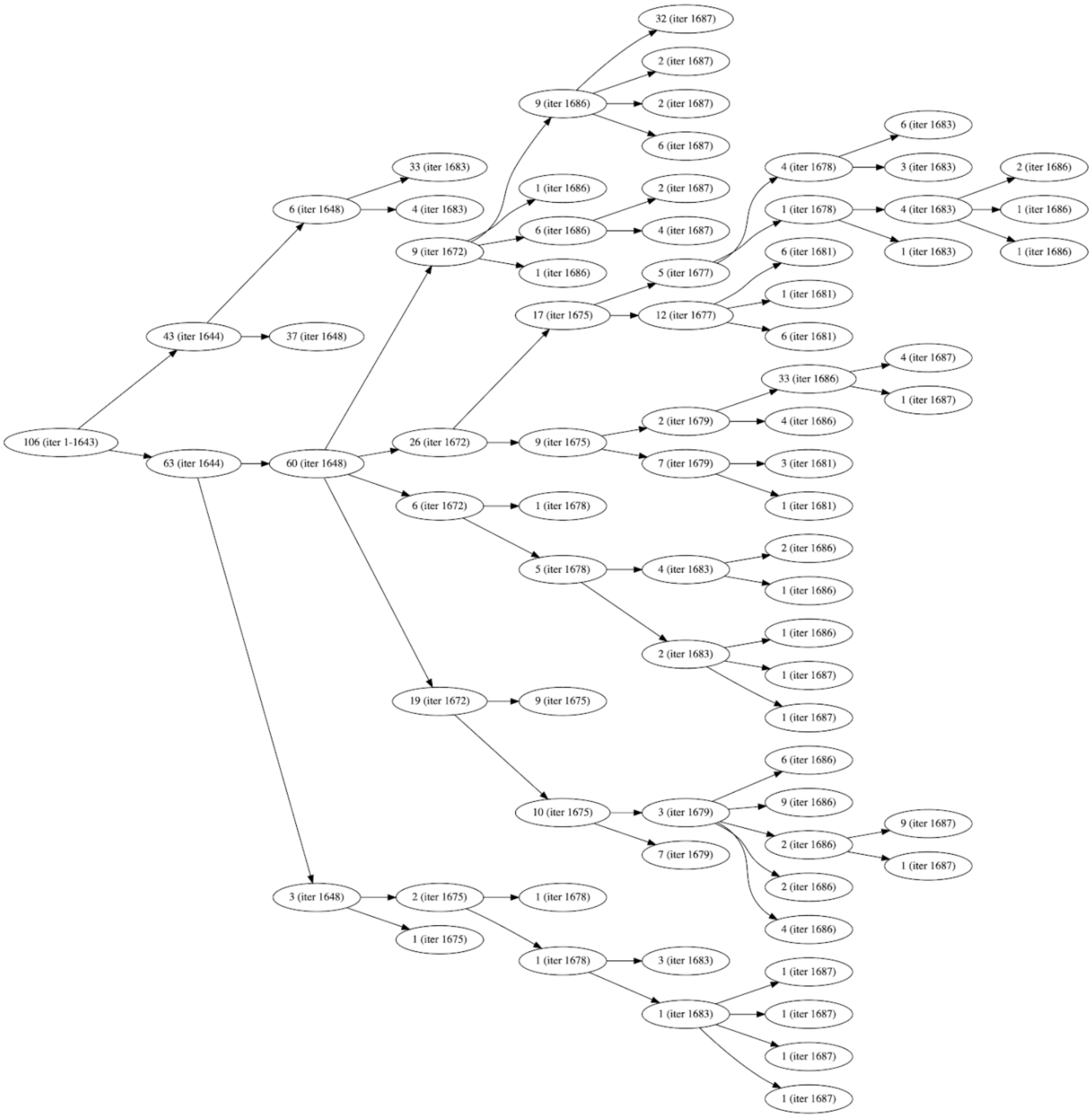

**Supplemental Fig. 12** Trajectory splitting tree of the 106 pathways that reached the "open" state. The number of each node indicates the number of pathways at the given WE iteration in parentheses. All trajectories shared the same parents until iteration 1643, the first splitting of trajectories occurring at iteration 1644. Note that the sum of the child pathways does not necessarily match up with the parent's number of pathways at subsequent iterations due to splitting and merging with other trajectories (not shown).

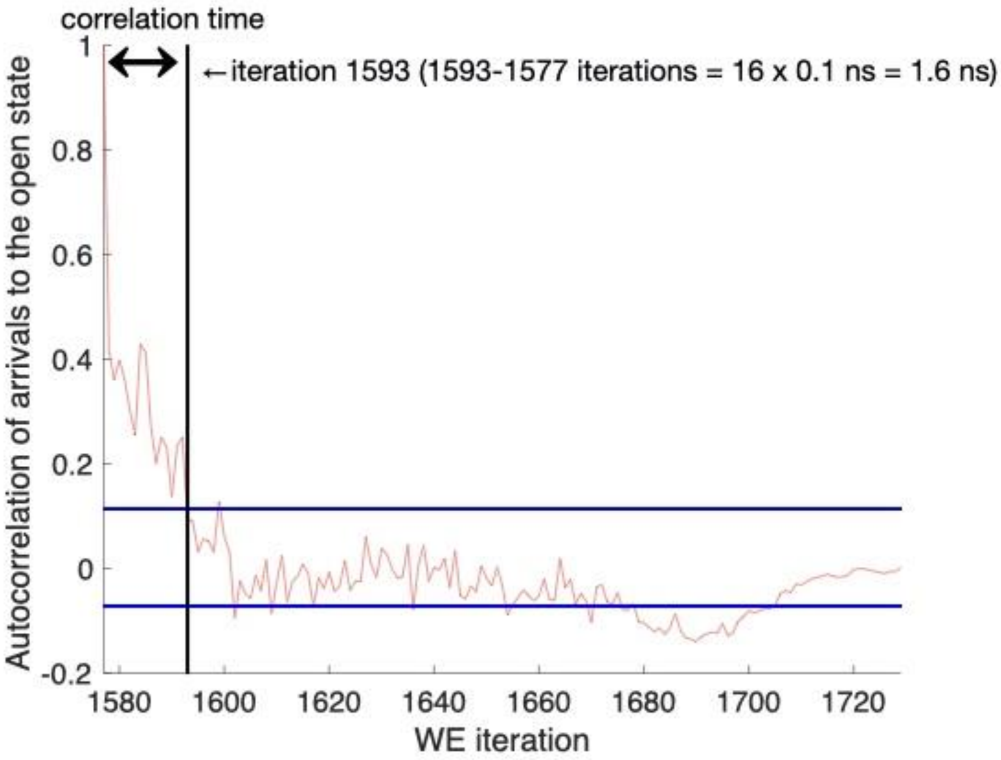

**Supplemental Fig. 13** Autocorrelation of arrivals from the “down” state to the “open” state (red) with a 95% confidence interval (blue). The confidence interval was generated using a Monte Carlo bootstrapping strategy where a bootstrap consisted of 1000 randomly drawn datasets (with replacement) from all “down”-to-“open” flux values. The vertical line marks the first point at which the autocorrelation falls within the confidence interval and is used to calculate the correlation time.

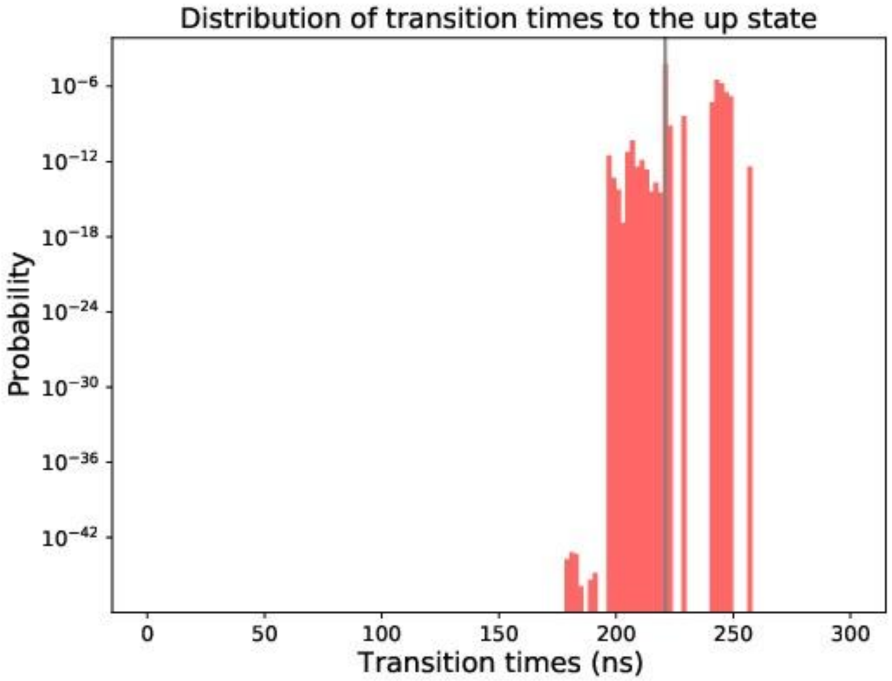

**Supplemental Fig. 14** Probability distribution of transition times from the “down” state to the “up” state. The most probable transition time is marked in grey. Note that the first 25% of the “fast” transitions are discarded here to calculate the most probable transition time.

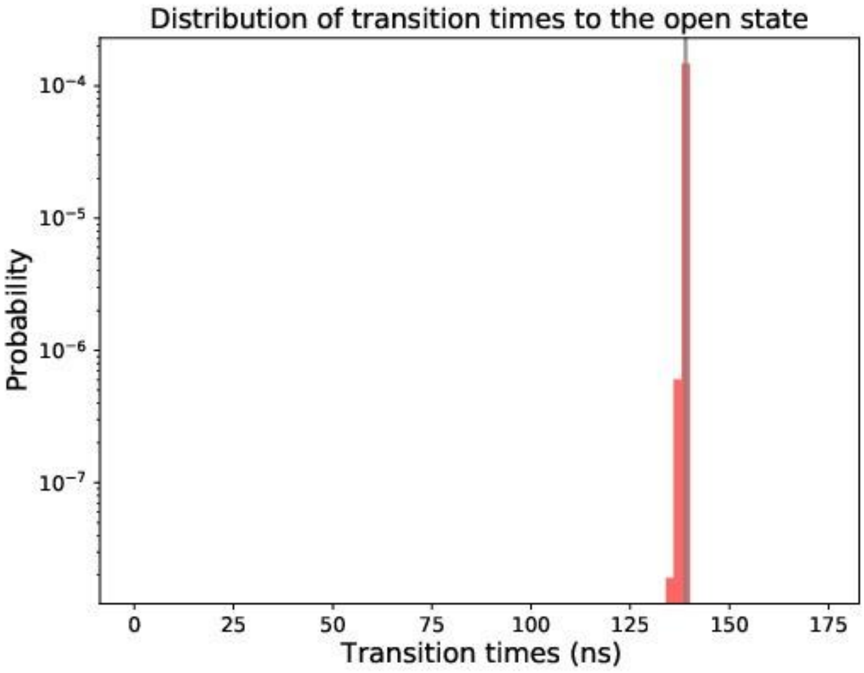

**Supplemental Fig. 15** Probability distribution of transition times from the "down" state to the "open" state. The most probable transition time is marked in grey. Note that the first 25% of the "fast" transitions are discarded here to calculate the most probable transition time.

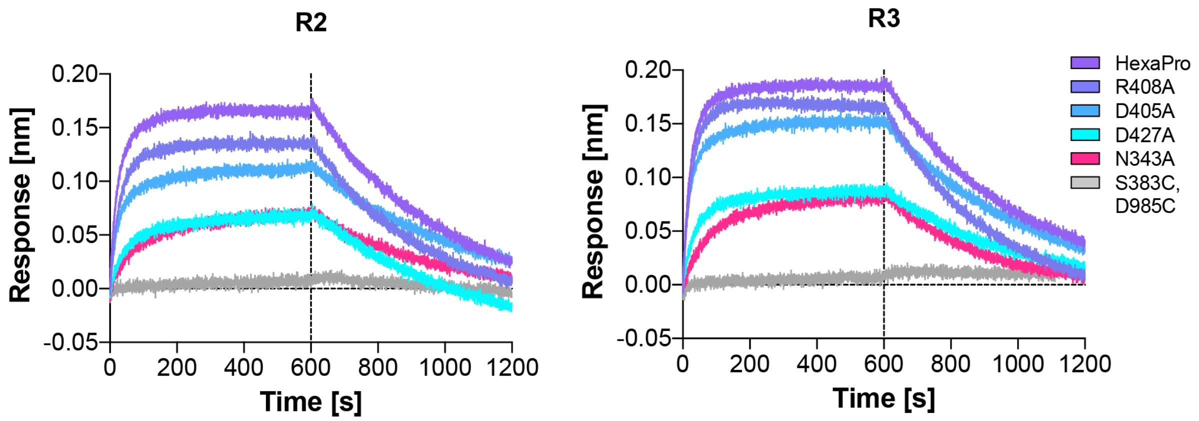

**Supplemental Fig. 16** BLI sensorgrams of spike variants binding to ACE2 from duplicate (R2) and triplicate (R3) experiments.

#### 3. Supplementary Tables

**Supplemental Table 1** Biolayer interferometry data of spike variants binding to ACE2.

| VARIANT | HEXAPRO | R408A | D405A | D427A | N343A |
|---|---|---|---|---|---|
| **R1 - Binding level (nm)** | 0.1733 | 0.1560 | 0.1206 | 0.0913 | 0.0783 |
| **R2 - Binding level (nm)** | 0.1776 | 0.1467 | 0.1208 | 0.0793 | 0.0751 |
| **R3 - Binding level (nm)** | 0.1831 | 0.1629 | 0.1506 | 0.0849 | 0.0816 |
| **Minimum (nm)** | 0.1733 | 0.1467 | 0.1206 | 0.07932 | 0.07512 |
| **Maximum (nm)** | 0.1831 | 0.1629 | 0.1506 | 0.09131 | 0.0816 |
| **Range (nm)** | 0.0098 | 0.0162 | 0.03 | 0.01199 | 0.00648 |
| **Mean (nm)** | 0.1780 | 0.1552 | 0.1307 | 0.0852 | 0.0783 |
| **Std. Deviation (± nm)** | 0.0049 | 0.0081 | 0.0173 | 0.0060 | 0.0032 |
| **Response (% to HexaPro)** | **100.00** | **87.19** | **73.43** | **47.85** | **44.01** |
| **Response decrease (%)** | **0.00** | **12.81** | **26.57** | **52.15** | **55.99** |

#### 4. Supplementary Videos

**Supplemental Video 1** Continuous pathway of RBD opening. This movie shows one of the continuous, unbiased pathways obtained from the WE simulations. All glycans are shown in blue except the N343 glycan which is colored magenta. Starting from all three RBDs in the "down" conformation, the chain A RBD lifts and twists counterclockwise into the "up" conformation, facilitated through interactions with the two adjacent RBDs, especially the N343 glycan gate on the chain B RBD. Upon reaching the "up" conformation, the RBD continues to twist into an "open" conformation en route to S1 dissociation.

**Supplemental Video 2** A comparison of the WE trajectory and ManifoldEM (MEM) CC1 and CC2 for a side view (PD 1386). It can be seen that there is strong agreement between the full WE trajectory and the sequential, piecewise combination of both CCs. Red arrows indicate direction of motion.

**Supplemental Video 3** A comparison of the WE trajectory and ManifoldEM (MEM) CC2 for a top-down view (PD 112). A strong agreement can be seen between the outputs of these two frameworks. To note, CC1 was not readily achievable from this view via manifold embedding, since the RBD-"down" to RBD-"up" trajectory from this view is orthogonal to the plane of the projection. Red arrows indicate direction of motion.

**Supplemental Video 4** Glycan gate at position N343 intercalates with residues to facilitate RBD opening. This movie zooms in closer to the glycan at position N343 to show how RBD opening is facilitated through intercalation between and underneath the residues F490, Y489, F456, F457 of RBD A. The glycan also transiently interacts with other residues of the RBD which are shown when they are within Å from the glycan.

**Supplemental Video 5** Mapping of residue contacts to RBD throughout opening pathway. Distances between residues throughout a continuous opening pathway calculated for the trajectory shown in **Supplemental Videos 1 and 2**. Distances to each residue from $RBD_A$ are shown for each chain in panels A-C and each of the glycans in panel D. Select regions are labeled, and N165, N234, and N343 are labeled with +, ++, +++, respectively.

(55) Liebschner, D.; Afonine, P. V.; Baker, M. L.; Bunkoczi, G.; Chen, V. B.; Croll, T. I.; Hintze, B.; Hung, L. W.; Jain, S.; McCoy, A. J.; Moriarty, N. W.; Oeffner, R. D.; Poon, B. K.; Prisant, M. G.; Read, R. J.; Richardson, J. S.; Richardson, D. C.; Sammito, M. D.; Sobolev, O. V.; Stockwell, D. H.; Terwilliger, T. C.; Urzhumtsev, A. G.; Videau, L. L.;

Williams, C. J.; Adams, P. D. Macromolecular Structure Determination Using X-Rays, Neutrons and Electrons: Recent Developments in Phenix. *Acta Crystallogr. Sect. D Struct. Biol.* **2019**, *75* (Pt 10), 861–877. https://doi.org/10.1107/S2059798319011471.

(56) Tang, G.; Peng, L.; Baldwin, P. R.; Mann, D. S.; Jiang, W.; Rees, I.; Ludtke, S. J. EMAN2: An Extensible Image Processing Suite for Electron Microscopy. *J. Struct. Biol.* **2007**, *157* (1), 38–46. https://doi.org/10.1016/j.jsb.2006.05.009.
